# Nanopore Decoding with Speed and Versatility for Data Storage

**DOI:** 10.1101/2024.06.18.599582

**Authors:** Kevin D. Volkel, Paul W. Hook, Albert Keung, Winston Timp, James M. Tuck

## Abstract

**Motivation:** As nanopore technology reaches ever higher throughput and accuracy, it becomes an increasingly viable candidate for reading out DNA data storage. Nanopore sequencing offers considerable flexibility by allowing long reads, real-time signal analysis, and the ability to read both DNA and RNA. We need flexible and efficient designs that match nanopore’s capabilities, but relatively few designs have been explored and many have significant inefficiency in read density, error rate, or compute time. To address these problems, we designed a new single-read per-strand decoder that achieves low byte error rates, offers high throughput, scales to long reads, and works well for both DNA and RNA molecules. We achieve these results through a novel soft decoding algorithm that can be effectively parallelized on a GPU. Our faster decoder allows us to study a wider range of system designs.

**Results:** We demonstrate our approach on HEDGES, a state-of-the-art DNA-constrained convolutional code. We implement one hard decoder that runs serially and two soft decoders that run on GPUs. Our evaluation for each decoder is applied to the same population of nanopore reads collected from a synthesized library of strands. These same strands are synthesized with a T7 promoter to enable RNA transcription and decoding. Our results show that the hard decoder has a byte error rate over 25%, while the prior state of the art soft decoder can achieve error rates of 2.25%. However, that design also suffers a low throughput of 183 seconds/read. Our new Alignment Matrix Trellis soft decoder improves throughput by 257x with the trade off of a higher byte error rate of 3.52% compared to the state-of-the-art. Furthermore, we use the faster speed of our algorithm to explore more design options. We show that read densities of 0.33 bits/base can be achieved, which is 4x larger than prior MSA-based decoders. We also compare RNA to DNA, and find that RNA has 85% as many error free reads as compared to DNA.

**Availability and implementation:** Source code for our soft decoder and data used to generate figures is available publicly in the Github repository https://github.com/dna-storage/hedges-soft-decoder (10.5281/zenodo.11454877). All raw FAST5/FASTQ data is available at 10.5281/zenodo.11985454 and 10.5281/zenodo.12014515.

## 1. Introduction

DNA has emerged as a viable data storage medium in recent years, with advancements focused on reducing synthesis costs (Nguyen et al. (2021)), improving encoding densities (Choi et al. (2019)), and selectively retrieving information from DNA libraries (Organick et al. (2018)). While many of the early works assumed high throughput sequencing technologies, the sequencing technology landscape has seen major changes due to the continual advancements in yield and accuracy in nanopore sequencing devices and their basecalling algorithms (Wang et al. (2021); Pag`es-Gallego and de Ridder (2023)). The ability to reach yields of 100 Gb per flow cell make nanopore sequencing a competitive option for large scale molecular storage systems in addition to portable ones (Yazdi et al. (2017)). Furthermore, nanopore sequencing enables long read sequencing, real time signal analysis (Kovaka et al. (2021)), and can directly interrogate other biopolymers such as RNA. Hence, nanopore sequencing has the potential to support a wide range of interesting storage system architectures, but few of these options have been deeply explored.

A current bottleneck for nanopore-based DNA storage systems are their high cost of decoding. Most studies rely on *post-hoc* MSA and clustering analyses as a critical decoding step. While some works may be able to write information at a density of 1.33 bits/base (Chen et al. (2021)), read density can be over an order of magnitude lower (0.079 bits/base) due to reading each base 16.8x times on average in order to build a consensus read that will correctly decode (Supplemental Figure 1). In the context of a storage system, this implies that the computational infrastructure supporting the decoding process and the sequencing material costs will be 16.8x larger than if each originally encoded strand was read once.

Convolutional codes (Press et al. (2020); Chandak et al. (2020)) have shown great promise for single-read approaches that can extract the information payload of a sequence from a single read. HEDGES (Press et al. (2020)) is a convolutional code that is tolerant of insertions and deletions and has been designed specifically for *single-read*, but it has only been evaluated for Illumina-based sequencing platforms, which have an order of magnitude lower error rate (0.1% in Tomek et al. (2019); Organick et al. (2018)) than to nanopore sequencing. HEDGES tolerates errors by systematically and serially guessing the location of errors, which can significantly increase compute time under the higher error rates of nanopore. Chandak *et al*. have shown that a *soft decoding* technique that directly integrates the base probabilities output by nanopore basecallers can substantially lower read costs and byte error rates. However, the decode throughput is low, 183 seconds per read on average based on our benchmarking measurements.

Building on the success of prior convolutional codes and basecaller integration (Chandak et al. (2020)), we explored two trellis soft decoders for the HEDGES encoding that can run in parallel on a GPU and solve for the most likely decoding. First, we integrate Chandak *et al*.’s trellis with the constrained encoding of HEDGES and parallelize it to run on a GPU. We use this as our baseline comparison. Second, we developed a new decoding algorithm that leverages a dynamic programming approach to compute trellis state probabilities in a way that can efficiently utilize the GPU’s parallelism and memory architecture.

In this work we perform a systematic comparison between hard decoding and each soft decoder. First, we use HEDGES on state of the art nanopore basecallers from Oxford Nanopore Technologies (ONT) and find that on average the byte error rate is 25.4% when sequencing 7 uniquely encoded and synthesized long DNA molecules of 2297 bp. This work provides the first study that has been done to directly compare soft and hard decoding performance of convolutional codes for the same population of nanopore reads. We show that Chandak *et al*.’s CTC decoding algorithm can significantly reduce byte error rate to 2.25% on average. However, because of the low throughput, we limited our analysis to just 2 encoded strands. We evaluated our new algorithm on this same sample of encodings and show that we provide comparable error rates (3.52%), but with a speedup of 257x compared to Chandak *et al*.’s soft decoder when evaluating both on GPU implementations. This speedup enables scaling to our full set of 7 strands to show that the error rate on this large sample is 2.59%. We then synthesized 10 additional strands spanning several lengths and encoding densities to understand the accuracy and density trade-offs. Based on our data, we project that our decoding can achieve read densities of 0.33 bits/base, 4× larger compared to coverage-optimized MSA approaches such as Chen et al. (2021) (Supplemental Figure 1). We also demonstrate our decoder’s flexibility by applying our algorithm to an open-source RNA basecaller, and we show that it achieves a lower byte error rate than DNA using the baseline HEDGES decoder with a state-of-the-art basecaller. This supports the feasibility of using RNA decoding as part of a data storage system.

## 2. Materials and Methods

### 2.1 Information Encoding

We employ the HEDGES code as our baseline in this work (Press et al. (2020)). We chose this encoding due to its ability to avoid repetitive bases, GC balancing constraints, and variable encoding densities. The HEDGES encoder builds a DNA strand based on the results of a hash algorithm that digests three pieces of information: *history bits, base index*, and the *next bit* to be encoded (Figure 1A). The history bits are used in conjunction with the base index to embed the context of each encoded bit within the base sequence. The approach of combining history information during encoding places HEDGES within the class of convolutional codes. Such codes are decoded by making a series of guesses about what information was stored. Thus, the hash and embedded context is designed to generate distinct DNA sequences that can be distinguished even in the instances of errors injected by the channel as guesses are generated.

**Fig. 1:**
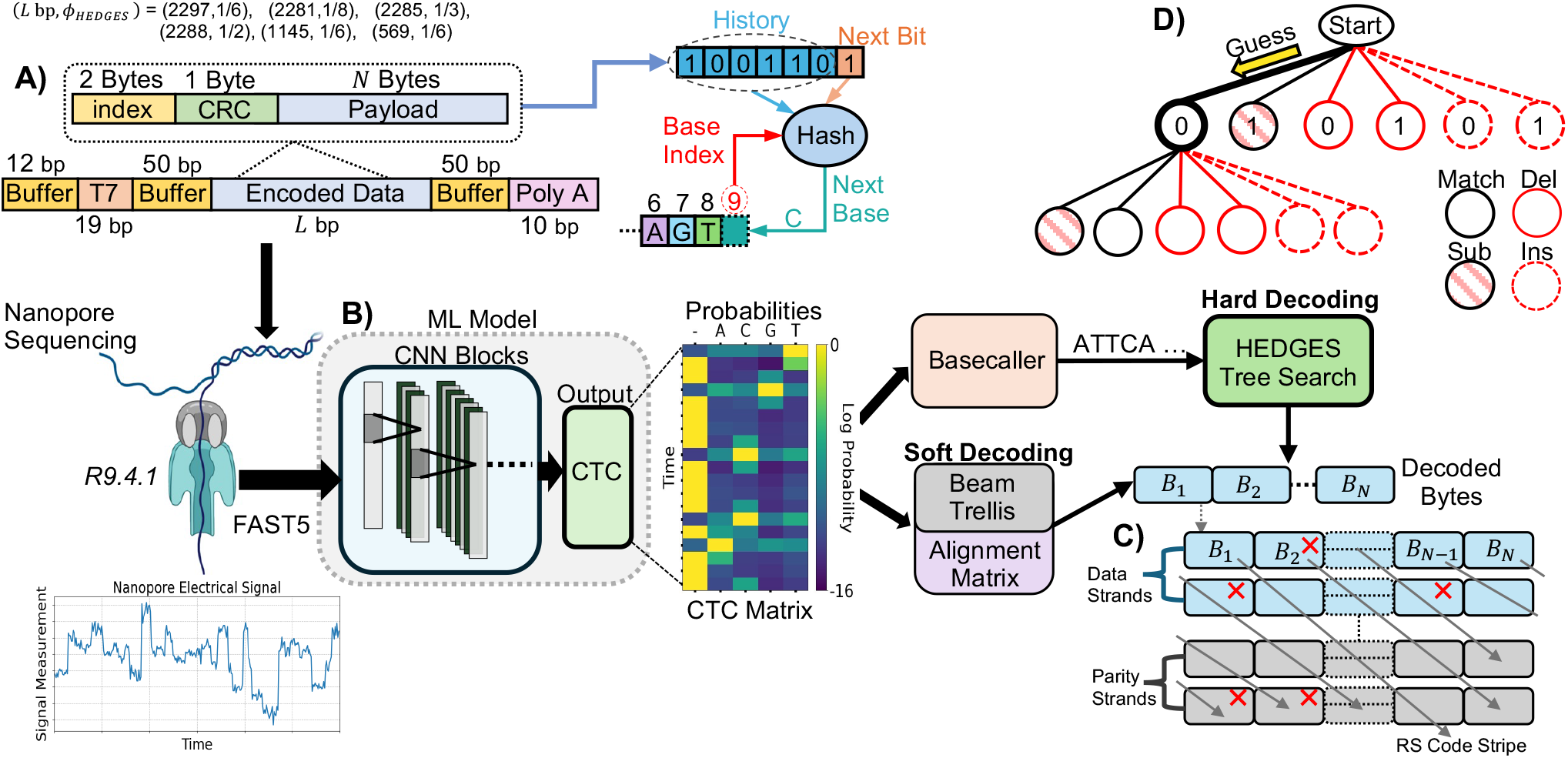
Overview of experimental workflow. **A)** Encoding parameters and strand design used throughout the course of our experiments. **B)** This work assumes a ML model that transforms nanopore signals to CTC outputs from which the code can be directly decoded (soft decoding), or decoded following a basecalling process that decodes the ML model output. **C)** Diagonally striped RS outer code model assumed to allow for final densities to be calculated when taking into account the rate of byte-errors emitted from the studied decoders. **D)** Outline of HEDGES decoding. Guesses are made on which bit was encoded which also emits a corresponding base according to the hash. Guesses include all possible error scenarios and are organized within a tree data structure.

### 2.2 Soft Decoding Algorithms

Soft decoders leverage probabilities associated with each symbol to decide the most likely message sent given the received symbol probabilities. Such decoders are widely used and well known to offer advantages over hard decoding. While most prior work in DNA storage relies on hard decoding of sequencer generated basecall data, nanopore sequencing workflows make it possible to extract detailed per base probabilities. Nanopore basecalling workflows typically consist of two main steps: using a machine learning (ML) model to generate scores for assignment of bases to the electrical signal, and interpreting the scores to produce a final sequence of bases (Figure 1B).

One ML model output commonly used for nanopore sequencing is the connectionist temporal classification (CTC) output (Neumann et al. (2022); Pag`es-Gallego and de Ridder (2023)). This output is formed as a matrix with two dimensions. One dimension being interpreted as time, and the other dimension corresponding to the alphabet that a message is constructed from. Each element of the matrix represents a log probability that a symbol of the alphabet or a blank occurs at a given CTC time step. Based on the same CTC data we consider two soft decoders that take different approaches to estimating message probabilities. A key insight for CTC model outputs is to be able to learn and tolerate time variation in symbol signals Graves et al. (2006). This has a natural application in nanopore sequencing considering dwell time variations that may occur as bases traverse the pore. Because of the time variation and probabilistic outputs, CTC outputs do not directly convey a single message. Instead, *CTC-encodings* are used to construct alignments of messages to the CTC data to calculate probabilities for the message. Such encodings allow for the representation of the same base symbol occupying multiple time steps, e.g. the encoded AAA decodes to a single base message A. However, to enable repeats in the decoded message, at least one blank (-) must be included to separate their CTC repeats from their decoded repeats. For example, the message AA is only allowed encodings of the form A-A.

The intuition behind both soft decoders in this work is to determine the message that best synchronizes with the CTC information by taking into account their different possible *CTC-encodings*. The soft decoder of Chandak *et al*. synchronizes messages by expanding the trellis complexity to evaluate message positions at every CTC time step. On the other hand, our approach considers and compares how well message prefixes align across all time steps, enabling a time and memory saving dynamic programming approach. As we will show, the approach taken to perform this synchronization significantly impacts the compute and memory complexity.

#### 2.2.1 Beam Trellis Algorithm

HEDGES was originally described with hard decoding, so we extend it to support soft decoding. Since it is a convolutional code, we construct a full *trellis* to represent its decoding steps. A trellis for HEDGES must have a width of at least 2^*H*^ states for *H* history bits. Traditionally, each state has 2 outgoing edges representing the transition to another history as new bits are added to a message. The next step is determining how to score each message with the CTC data while accounting its various *CTC-encoding* alignments.

The approach of Chandak et al. (2020) to incorporating *CTC-encodings* into a trellis is to extend the number of states by a factor of the length of the encoded strand (*L*) for a total of 2^*H*^ *L* states (Figure 2A). Now, each state represents a value of history at a given message index. In this approach, each state is updated a number of times equal to the time dimension of the CTC matrix (*T*). During the updating process for some state at trellis-step *t* + 1 three candidates are considered from the previous step *t*. Two candidates advance the index of the decoded strand (*S*_*X*_, *S*_*Y*_), while the remaining *S*_*W*_ does not (Figure 2A). The state *S*_*W*_ is the mechanism by which *CTC-encodings* that allow for a symbol to occupy multiple time steps are accounted for in decoding a fixed length message. Thus, every state carries a *non-blank* and *blank* score portion, which are combined together when calculating the total score for a transition (Figure 2B). With multiple states advancing the decoded strand index differently, edges representing *CTC-encodings* that convey the same message may occur which requires that they are merged so that an accurate score for a message can be obtained (Figure 2C). Given this algorithm’s similarities to so-called Beam search algorithms (Scheidl et al. (2018)), we refer to this approach as the Beam Trellis.

**Fig. 2:**
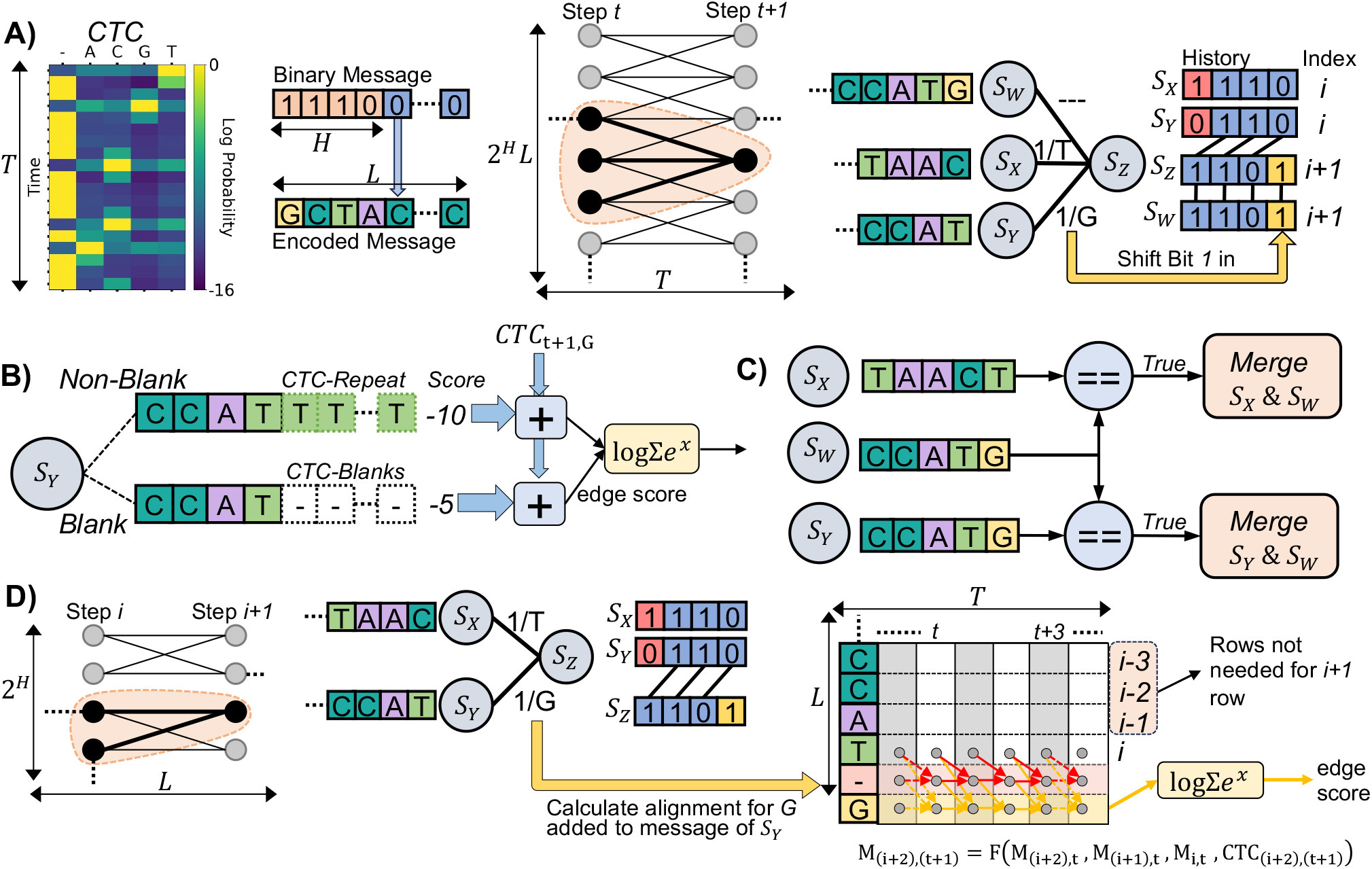
Soft decoding algorithms evaluated in this work. **A)** The trellis architecture and connections of states used by the Beam Trellis algorithm (Chandak *et al*.). **B)** Edge scoring mechanism of the Beam Trellis algorithm to account for *CTC-encodings* of candidate messages. **C)** State merging required by the Beam Trellis algorithm to account for duplicate state messages. **D)** Architecture and scoring methodology used by our novel Alignment Matrix algorithm. Arrows in the Alignment Matrix represent data used to calculate newly added rows. Red (yellow) arrows indicate the data dependencies for the added blank (new base) symbol.

The time complexity of this algorithm is *O*(*L*^2^*T* 2^*H*^) because of the *L*2^*H*^ number of states that are evaluated *T* times. The additional factor of *L* arises from the need to compare incoming messages of length *L* when evaluating each incoming edge for a state to determine if there are multiple *CTC-encodings* representing the same message. Likewise, since each state must store a complete message of length *L*, space complexity can be written as *O*(2^*H*^ *L*^2^). Provided that *T* will grow proportionally to *L, T* factors can be replaced by *L* in the complexity expressions. From this, we can see that this algorithm has poor time and memory scaling as the length of messages increase.

#### 2.2.2 Alignment Matrix Algorithm

Our novel approach to integrating CTC information into a trellis is to calculate the alignment for each message of the 2^*H*^ states directly (Figure 2D). This is done by using the algorithm of Graves et al. (2006) to calculate the so called forward variables that represent the total probability of a prefix for a message for a certain time step. Conceptually, each forward variable for a base in a message is stored in a matrix *M* of dimension *L* × *T*. Each element of *M* (*i, t*) representing the sum of probabilities of all *CTC-encodings* of *L*[0 : *i*], the prefix of *L* up to and including position *i* for some time *t*. Thus, when transitioning from decoder step *i* to *i* + 1 we can calculate *M* (*i* + 1, *t*) for all *t*, e.g row *i* + 1 of *M*. Because this row represents the probability of each newly constructed message prefix at each time step, it can be used as a method to score each transition edge in the trellis. To transform the row of values into a scalar for comparison, we compute the log sum of exponentials over the log probabilities of the freshly computed row. Our reasoning for using this value is that it represents a total probability for a prefix across all times *t*.

In practice, to account for paths to a base that pass through blank symbols, a row for a blank is included in *M* previous to the newly added base as shown in Figure 2D. Each of these newly added rows (blank and non-blank) can be completely derived from the most recent row corresponding to a base in the message. Thus, we do not store the entire matrix for every state or recalculate it every moment that it is needed. Instead we only store the most recently calculated row, and only perform computations needed to calculate the new row for a newly added base. In concrete terms, the row related to a base with blanks included is calculated as 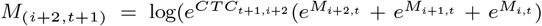. What this calculation represents is a summation of all probabilities for paths into the new base while accounting for the probability of the new base at the given time step as determined by the CTC matrix. This calculation based on Figure 2D assumes that the base added by a trellis edge is different than the most recent base (*G* ≠ *T*). If this were not the case, the term 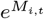 is removed from the calculation because at least one blank character is needed as previously discussed to encode message repeats.

The time complexity of our algorithm is *O*(2^*H*^ *LT*). We determine this given that the trellis has 2^*H*^ states and *L* trellis evaluation steps. At each evaluation, we must calculate an alignment across *O*(*T*) time steps for each edge incoming to each state. Assuming the number of edges and the number of bases added to a message on each transition is fixed for a given code we assume these factors to be *O*(1). Thus, the time complexity of a single trellis propagation step is 2^*H*^ *T*, and the complexity of *O*(2^*H*^ *LT*) follows. This complexity is made possible by the dynamic programming approach taken to calculating rows for new bases, rather than re-calculating rows already visited.

Memory complexity follows a similar argument, where we only store 1 row of a matrix for each state so that alignments can propagate. Thus, a memory complexity of *O*(2^*H*^ *T*) is achieved for our algorithm. Assuming *T* is proportional to *L*, our algorithm reduces complexity by a factor of *L*. A detailed pseudo code description of the Alignment Matrix algorithm can be found in Supplemental Section E.

#### 2.2.3 GPU Parallelization

We recognize that to perform all of the necessary calculations across each trellis approach, significant computational effort will be required to aligning long messages with their correspondingly long CTC matrix. Provided the abundance of independent calculations that can be performed across states of the Beam Trellis algorithm and the rows of the matrix in the Alignment Matrix algorithm, we leverage GPUs to accelerate each algorithm. In our implementations we strive to utilize best practices by considering occupancy, shared memory resources, and memory access coalescing patterns. Details of our GPU implementations for both soft decoders can be found in Supplemental Section A.9.

When benchmarking soft decoder algorithms they are run on nodes consisting of a NVIDIA RTX 2060 Super GPU device, a single AMD EPYC 7302P 16-Core processor, and 128GB DDR4 DRAM. The baseline HEDGES decoder runs on either Intel Xeon Gold 6226 or 6130 processors that have 192 GB RAM per node.

### 2.3 Experiment Workflow

#### 2.3.1 Encoding Information in Molecules

To complete our evaluation of decoders, we encode and synthesize 17 unique DNA template molecules following Figure 1A. A primary goal for our analysis is to understand how information error rates will be influenced by the rate of the encoding and the length of a molecule passing through a pore. Thus, our designs cover HEDGES code rates of 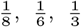, and 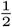. Each encoded strand was bookended by signal buffer sequences of length 50 bp to protect information carrying bases from transient behaviour entering and leaving the nanopore. On the 3^′^ end we also include an additional 10 base poly-A tail, and on the 5^′^ end we allocate 19 bp for a T7 promoter to allow for transcription of RNA molecules and an additional 8 bp for a synthesis buffer. For every design, we keep these additional 5^′^ and 3^′^ bases constant. The full length of the 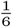 rate strands with additional regions is 2297 bp. For the rates of 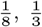, and 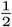 we aim to keep strand length relatively constant with their respective entire lengths being 2281, 2285, and 2288 bp. For the ^1^ rate we synthesize 2 additional length of molecules that are 1145 bp and 569 bp to study short length strand impacts on nanopore sequencing. For HEDGES parameterization, we limit homopolymers to be a maximum of 3 and fix a GC content of 50% over 12 base pair windows. Each strand was ordered as DNA gBlocks Gene Fragments from Integrated DNA Technologies.

Our molecules are derived from several sources of data. The strands for the 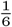 HEDGES rate and 2297 bp design are all derived from the same thumbnail image of the periodic table symbol for phosphorous. The 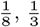, and 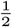 HEDGES rates strands encode the complete 8th, 4th, and 6th amendments of the Constitution of the United States respectively. The 1145 and 569 bp with 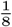 HEDGES rate designs encode the first 76 characters and characters 28 to 56 of the 4th amendment respectively. Copies of the raw encoded data and exact strands synthesized for these experiments are included within our public code release.

#### 2.3.2 Nanopore Sequencing and Preprocessing

With our synthesized DNA molecules, we sequence all strands using ONT nanopores of version R9.4.1. For each sequencing run, we use the latest available ONT basecalling models at the time of sequencing as reported in Supplemental Table 1 to generate FASTQ information. We used this initial FASTQ data in order to demultiplex individual reads to their original encoded strand and to eliminate reads that we do not want to impact measured decode rates from our decoders. Demultiplexing is done via the encoded index and CRC bytes within the encoded strand (Figure 1A). Further details are available in Extended Methods A.

Using HEDGES to decode basecalls, we attribute each read to an encoded strand if the decoded index bytes is in agreement with the CRC byte. While performing read attribution, we eliminate short reads and exceedingly long chimeric reads (Supplemental Figure 2). Our reason to exclude these reads is to obtain a clearer understanding of decoder performance on the characteristic of nanopore signals and not information loss that may occur from small fragments that result from other molecular handling and processing steps. We verify that our preprocessing does not bias the quality of reads to significantly higher qualities, and eliminated reads correlate to outlier low quality reads that would be considered failed reads by ONT (Q Score *<* 9). When considering reads of Q Score *>* 9 Supplemental Figure 5 shows when demultiplexing a 5 strand sequencing run of 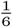 rate 2297 bp strands the average Q Score of retained reads varies between 13.37 − 13.57 compared to 13.25 of the entire sequencing run.

To analyze decoding accuracy performance and basecall error rates while limiting computational overhead we randomly sample our set of attributed reads to sizes appropriate for our analysis needs. To ensure high confidence in basecall error rates and error patterns, we take a subset of 100k reads for the baseline hard decoder. We analyze decoder performance and basecall error patterns for the chosen 100k subset against the original ONT basecall FASTQ. We also collect the FAST5 data corresponding to this subset so that basecall information of the CTC model used for soft decoding can be analyzed (Figure 1B). We use the latest CTC version of the open source ONT Bonito basecaller available, and the exact code version of this model is included in our code release repository. For soft decoding, we are only interested in byte error rate and so we reduce the number of samples. For our novel soft decoder, we take a 10k subset of reads from the original 100k subset for each strand. In our evaluation of Chandak et al. (2020)’s CTC decoder we experienced slow throughputs caused by the complexity of the decoder. Thus, we use a 2k subset sampled randomly from the 10k set.

#### 2.3.3 CTC Data Orientation and Buffer Regions

CTC matrices received from the ML model will not only include data related to the payload, but also buffer regions. Leaving data within the CTC matrix related to to buffer regions can potentially disrupt the soft decoding algorithms by causing alignments of payload bases to signals that are unrelated. Furthermore, DNA molecules can be sequenced as their encoded forward versions or their reverse-complement. This is important to consider so that the decoder can generate the appropriate bases in the trellis. To solve this problem with only CTC information, we adapt the techniques of Kürzinger et al. (2020) to locate and trim buffer region CTC data (Supplemental Section A.8).

#### 2.3.4 Measuring Byte Error Rate and System Density

For all bytes within each strand, we calculate the byte error rate as the total number of decoding failures observed for each individual byte. To account for biasing that occurs in HEDGES decoding where bytes towards the end of a strand have a higher failure rate (Supplemental Figure 12), we calculate a mean rate 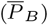 for byte errors across all positions and encoded strands (Supplemental Equation 3). Following the methodology if Press et al. (2020), we use this average byte error rate to model the effective error rate that is observed in a Reed Solomon (RS) codeword with diagonally striped bytes between strands (Figure 1C). Using 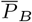, we calculate the probability to decode a RS codeword (Supplemental Equation 10) and choose a design that can access 1 TB with a mean time to failure (MTTF) of 10^6^ accesses. With the outer code design, we calculate a complete density, *ϕ*, in bits per base following the steps of Supplemental Section A.5.

## 3. Results

### 3.1 Hard Decoding Byte Errors

When considering the baseline hard decoder of the HEDGES code we want to understand both the rate bytes can be recovered and also the computational effort expended for a given byte error rate. As shown in Figure 1D, the HEDGES decoding algorithm forms a tree representing guesses that can be made on what information was encoded and what errors may have occurred within the basecalled sequence. Figure 3 demonstrates the relationship of compute time and byte error rate for the two basecallers considered where *DNA-CTC* is the CTC output based model used as the basis of our soft decoder algorithms and *DNA-ONT* represents ONT production basecallers (Supplemental Table 1). Both basecallers are applied to the same subset of 100k reads for all 7 strands of the 2297 bp 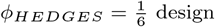.

**Fig. 3:**
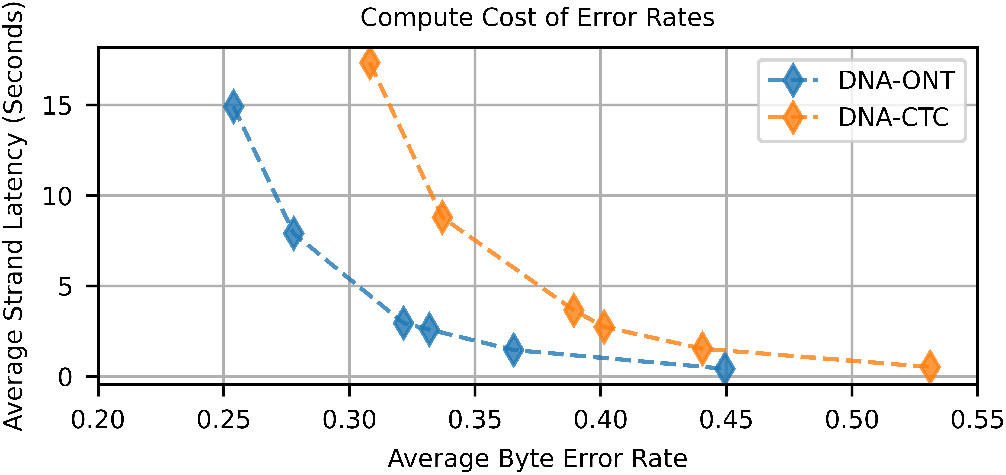
Average strand decode time versus average byte error rate when using the HEDGES baseline hard decoding algorithm for ONT and CTC basecalling models.

We find that for both basecallers there are diminishing returns when allocating more guesses to the algorithm indicating that it is intractable to reach low byte error rates by simply increasing compute effort. The lowest mean byte error rates achieved for the ONT and CTC models were measured to be 0.254 and 0.308 respectively when allowing a limit of 6 million guesses. When placing these byte error rates in our approach to determining overall system density, we find that hard decoding error rates cannot meet our target MTTF = 10^6^ for systems larger than 1 TB (Supplemental Figure 13). The higher byte error rate of decoding the CTC basecaller also serves as a control to show that the CTC model does not provide an unfair advantage in the quality of information produced by the ML model compared to the ONT models. Corresponding to this higher byte error rate, we find that CTC basecaller base error rates are 7.6% on average and are higher than the ONT basecallers base error rates (5.63%, Supplemental Figures 3 and 4).

**Fig. 4:**
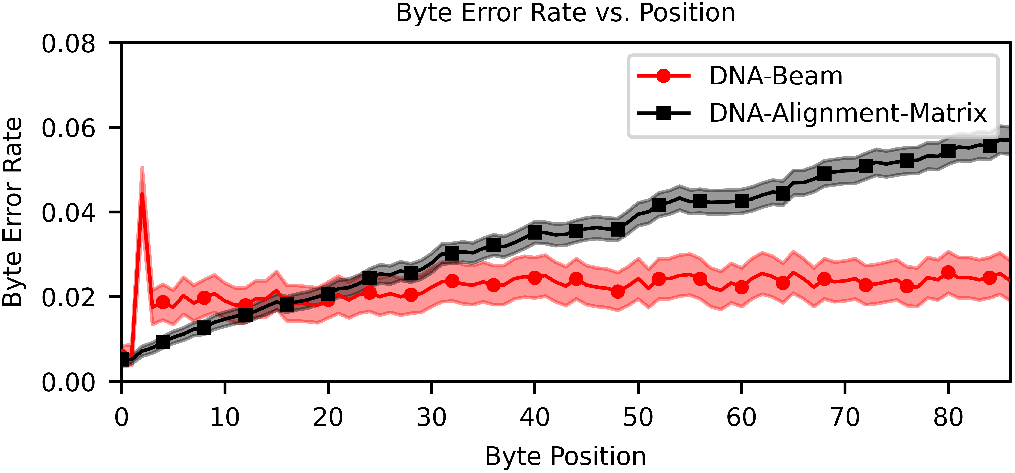
Byte error rate versus byte position for both soft decoders. Points represent mean error rate across encoded strands (Supplemental Equation 2), error bars calculated with Supplemental Equation 5 estimate a 95% confidence interval.

### 3.2 Soft Decoding Performance Analysis

Table 1 reports key metrics when comparing all three decoding algorithms. Error rates and densities in this table are derived from a subset of 2 of the 7 2297 bp 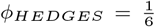 design strands. Our analysis demonstrates that for the same encoded strands, the Beam Trellis algorithm can reduce 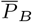 by an order of magnitude compared to hard decoding using 6 million guesses. We also find that our novel Alignment Matrix algorithm greatly improves 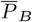 on this data set to 3.52%. The slight increase in byte error rate is a product of the error profile of byte errors within a strand as shown in Figure 4. We find that the byte error rate of the Beam Trellis algorithm remains relatively constant across the length of the strand, while the bytes at the end of a strand have higher error rates when decoded by the Alignment Matrix algorithm. This is caused by error cascades that are generated by the prefix probability scoring metric from data dependencies in the matrix we use to store alignments in Figure 2D. If a base is chosen such that it negatively impacts the score of successive bases that represent the correct path through a trellis, then it can become difficult to build a high enough score to correct wrong path choices when only a few bases at most are added to messages between comparisons (Supplemental Figure 9). On the other hand, the Beam Trellis algorithm has the flexibility to adjust states representing message indexes to best align to the CTC matrix.

**Table 1.** Comparison of decoding metrics for algorithm and molecule combinations. *T*_read_ measures just decode time excluding CTC model time. All hard decoders use ONT basecaller models.

| Molecule | Algorithm | $T_{read}$<br>s/read | $\bar{P}_B$ | $\phi$<br>bits/bp | Error<br>Free<br>Reads |
| --- | --- | --- | --- | --- | --- |
| DNA | Alignment Matrix | 0.713 | 3.52% | 0.203 | 87.15% |
|  | Beam | 183.17 | 2.25% | 0.221 | 84.08% |
|  | Tree-6M (hard) | 12.78 | 23.44% | 0.0128 | 58% |
|  | Tree-1M (hard) | 2.7 | 29.74% | N/A | 49.5% |
| RNA | Alignment Matrix | 1.06 | 12.52% | 0.106 | 74.34% |
|  | Tree-6M (hard) | 16.29 | 23.61% | 0.0128 | 56.72% |
|  | Tree-1M (hard) | 3.46 | 32.62% | N/A | 43.72% |

While the Beam Trellis algorithm does decrease 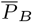 by 39% compared to the Alignment Matrix algorithm, in practice the impact on density is quite small (8.8% increase). Furthermore, the Alignment Matrix algorithm has a 257x larger throughput when we benchmark on a direct comparison of decoding 400 CTC matrices that we extracted from their respective reads. Our large gains in performance are derived from 2 factors. One, the computational complexity of our algorithm is reduced by a factor of *L*, and given we are decoding long messages, this can significantly impact the practical performance of the algorithms. Second, the limited complexity of our trellis architecture enables batching multiple reads to increase GPU utilization (Figure 5).

**Fig. 5:**
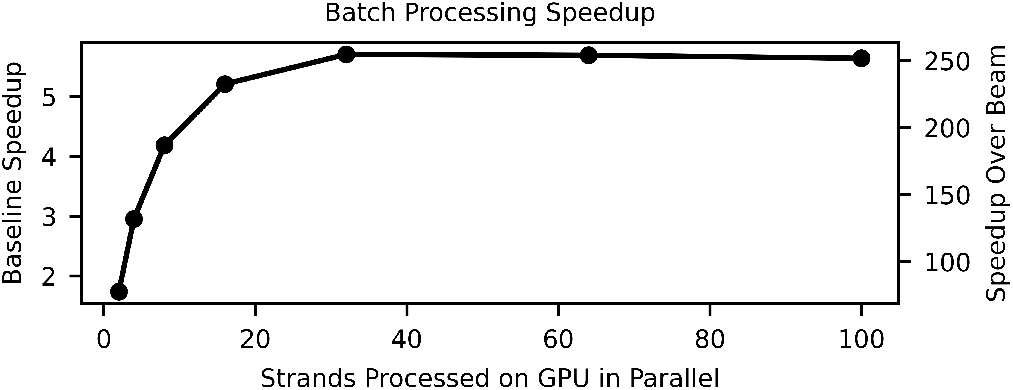
Speedups achieved with Alignment Matrix algorithm when batching multiple reads for GPU computation compared to decoding a single read (left y-axis) and compared to the decode rate of Beam Trellis algorithm (right y-axis).

For this strand design, the Alignment Matrix algorithm launches 3072 threads per read (Supplemental Figure 9). Given the total thread occupancy of our GPUs 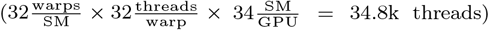, latency can be hid between data dependencies in the matrix by increasing the number of parallel reads decoding. However, for the Beam Trellis algorithm, we do not consider batching given that we launch approximately 552k threads per read which already saturates GPU thread occupancy (Supplemental Figure 8).

We also perform an analytic estimation of the total memory footprint of the two soft decoders (Supplemental Figure 10). We found that for the 400 reads used for benchmarking, the amount of memory used for each read on average is 52x lower when using the Alignment Matrix algorithm (0.032 GB) versus the Beam Trellis algorithm (1.66 GB).

Using the same 2 strands, we demonstrate the versatility of soft decoding CTC data by applying our decoder library with no changes directly to the CTC outputs of the open source RODAN RNA basecaller Neumann et al. (2022). Table 1 shows that again, CTC soft decoding outperforms hard decoding even when using ONT production basecallers. While we do find RNA has about 15% less total error free reads compared to DNA soft decoding, these results imply that storage systems that rely on RNA are feasible even under the conservative single-read assumption.

### 3.3 Optimizing Alignment Matrix Parameters

Given the computational advancements made by Alignment Matrix algorithm it is now possible to evaluate and understand parameter choices for the decoder. The parameters that we consider are strand length, HEDGES encoding density, and the reads chosen to provide to the decoder. We choose strand length to understand how the computational complexity of the decoder impacts decode time in practice. Also, given the positional dependence on error rate for bytes, we are interested in understanding how this profile may change with shorter strands. When changing the encoding rate we want to understand the density tradeoff of higher density encodings with the corresponding byte error rate that must be designed around. Lastly, we consider if information density can be significantly influenced by the quality of reads given to the decoder.

Table 2 summarizes decode throughput, 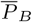, and *ϕ* for each design and when we consider reads of all quality in our data set (Nominal) and when reads are chosen from the set of reads with Q Scores in the range [15.1, 15.4]. Decoding rate as a function of strand length shows that the rate of information decoded outpaces the rate of information lost from a strand when shortening strands. For example, the 569 bp, 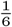 design has 383 bases of information after removing buffers and indexing and decodes at 0.0719 seconds read, while the 2297bp, 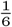 design encodes information over 2112bp and decodes at 0.723 seconds/read. So, while the shorter strand needs 5.5x as many strands to encode the same information, we can infer that decoding this total amount of information is 1.8x faster. Comparing 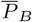 for shorter strands shows that when there are less bytes within a strand the overall byte error decreases by reducing impacts of cascaded errors (Supplemental Figure 15). However, the overall density (*ϕ*) is largest for the 2297 bp, 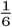 design because it can amortize overhead related to indexing and buffer regions more efficiently. These results indicate that strand length may be tuned to maximize density or byte throughput.

**Table 2.** Table of timing benchmarking, byte error rate 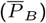, and density for 6 strand designs that are encoded and synthesized for experimental evaluation. *Nominal* refers to the case where each strand in each design is evaluated over a 10k read set derived from the entire space of viable sequencing reads. *Q Score* refers to analysis done when considering a subset of 10k reads that all have Q Scores in the range of 15.1 and 15.4. Seconds/read includes both ML model and decoder time, and decoding is done with a batch size of 50.

| Design<br>$L(\text{bp}), \phi_{\text{HEDGES}}$ | $T_{\text{read}}$<br>s/read | Nominal | | Q Score<br>[15.1, 15.4] | |
| --- | --- | --- | --- | --- | --- |
| | | $\bar{P}_B$ | $\phi$<br>bits/bp | $\bar{P}_B$ | $\phi$<br>bits/bp |
| 2281, 1/8 | 0.645 | 1.27% | 0.175 | 0.92% | 0.181 |
| 2297, 1/6 | 0.723 | 2.59% | 0.215 | 1.12% | 0.241 |
| 1145, 1/6 | 0.209 | 1.77% | 0.203 | 0.93% | 0.218 |
| 569, 1/6 | 0.0719 | 0.81% | 0.163 | 0.29% | 0.174 |
| 2285, 1/3 | 0.761 | 22.55% | 0.041 | 10.17% | 0.262 |
| 2288, 1/2 | 0.82 | 56.34% | n/a | 37.18% | n/a |

When comparing changes in *ϕ*_HEDGES_, we find that between any two encoding densities the lower density has a lower 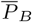. However, this does not always lead to higher densities. For example, 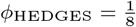 has 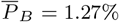 and 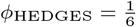 has 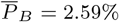, but the density for the former is 0.175 bits/bp while 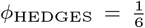 can achieve a 0.215 bits/bp with its error rate. However, 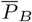 becomes too large to build efficient RS codes for the remaining higher encoding densities of 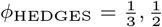. We consider if passing better quality reads can increase the viability of larger *ϕ*_HEDGES_ by evaluating 10k reads in 15 Q-score bins ranging from 10.9 to 15.1. By controlling the Q Score, we show that 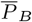 can be reduced greatly for 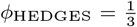 from 27% for reads in the Q score range [10.9 − 11.2] to 10.17% for reads in the range of [15.1 − 15.4] (Supplemental Figure 16). This leads to a 6.4x increase in density for 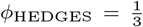 when higher quality reads are given to the decoder (Table 2).

We combine our findings together in Figure 6 by tuning strand length for each code rate when assuming error rates are for Q scores in the range of [15.1 − 15.4]. This allows for projections to be made about the maximum density that can be achieved with our decoder. In this analysis we make the assumption that the measured error rate versus byte position can be truncated to emulate shorter strand lengths than what were synthesized. This analysis shows the largest *ϕ* = 0.33 bits/bp is reached for 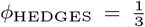 and strand length of 950 bp. Compared to the highest read density of Supplemental Figure 1 (Chen et al. (2021)), this projected density is 4x larger. These curves also indicate that limiting 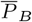 with shorter strands for larger encoding densities is important to maximize density, but for 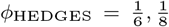 their 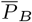 is low enough such that longer strands are preferred in order to amortize the overhead of bases associated with indexing or overhead for functional sites.

**Fig. 6:**
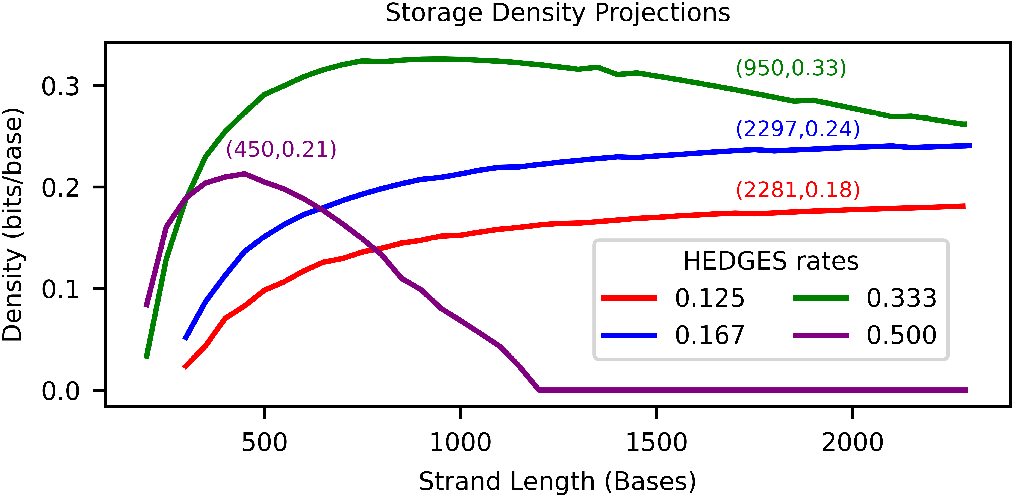
Projected densities when optimizing parameters of the Alignment Matrix algorithm.

## 4. Conclusion

Most DNA storage systems remain at a scale less than 1 GB and can tolerate slow decoding. However, to scale to large capacities and to advance our understanding of these systems, decoders must improve their throughput, read density, and support for varying strand length, molecule type, and encoding density. Our decoder is versatile and a step in that direction, but not the final step.

Further improvements in speed and error rate are needed and highly possible. Porting to more powerful GPUs will deliver speedups proportional to their threading capacity since Alignment Matrix is largely compute-bound and has only modest memory needs. Soft decoding is a key reason for low error rates, and we demonstrated Alignment Matrix works on outputs from both Bonito and RODAN on DNA and RNA, respectively. This implies that our approach is able to work independently of a particular model. We expect that our approach can benefit from advances in basecaller models as they are released. However, to ensure they remain compatible in the long run, it may be important to adapt to other common model outputs such as conditional random fields (Pag`es-Gallego and de Ridder (2023)). Additionally, the learned model could be coupled with the codeword space by leveraging application-specific training to improve inference quality of strands specific to the encoding (Wick et al. (2019)).

## A Extended Methods

### A.1 MSA Analysis

Supplemental Figure 1 provides an overview of read and write densities that are achieved by prior works that use MSA for decoding nanopore sequencing reads. We calculate read density by considering both the write density and the sequencing coverage used that each point is labeled with. Given that the write density is the base rate in which information is represented in terms of bases, and coverage represents on average the number copies of each encoded base, we then calculate read density as simply write density*/*coverage.

**Fig. 1:**
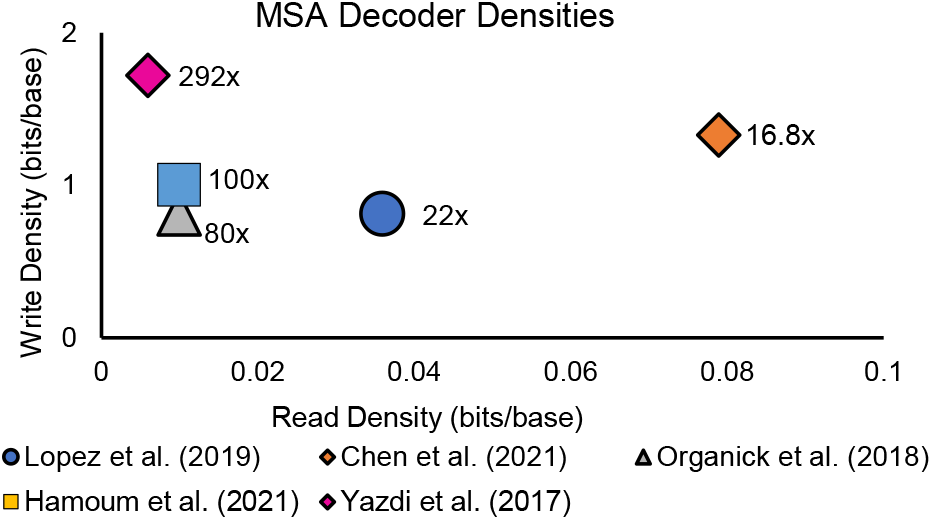
Read/write information densities of MSA decoders used for nanopore sequencing.

### A.2 Synthesis, Sequencing, and Transcription

#### A.2.1 Ordering and reconstitution

All designed strands were ordered as DNA gBlocks Gene Fragments from Integrated DNA Technologies (IDT). Upon arrival, dried fragments were reconstituted in either nuclease-free water or IDTE pH 8.0 (IDT, 11-05-01-13) and incubated at 50°C for 20 minutes. Reconstituted DNA fragments were stored at 4°C until use.

#### A.2.2 *In vitro* transcription (IVT)

IVT was performed for each strand separately with HiScribe T7 Quick High Yield RNA Synthesis Kit (NEB, E2050) according to manufacturer’s recommendations. Briefly, DNA fragments (70-100 ng) were mixed with 10 *µ*L of NTP Buffer Mix, 2 *µ*L T7 RNA Polymerase Mix, and nuclease-free water. In some reactions, 1 uL of 100 mM DTT was added due to a change in the manufacturer’s protocol. Reactions were then incubated at 37°C for 2 hours. Next, template DNA was removed by adding 30 *µ*L of nuclease-free water and 2 *µ*L DNase I (NEB, M0303S) to each reaction followed by incubation at 37°C for 15 minutes. RNA was then purified using the Monarch RNA Cleanup Kit (NEB, T2050L) following manufacturer’s recommendations. RNA was eluted in 50 *µ*L nuclease-free water and yield was quantified using Qubit RNA high sensitivity assay (Thermo Fisher Scientific, Q32852). RNA size was quantified using either RNA ScreenTape (Agilent, 5067-5576) or high sensitivity RNA ScreenTape (Agilent, 5067-5579) run on a TapeStation 4200 (Agilent, G2991BA). RNA was stored at -80°C until sequencing.

#### A.2.3. Nanopore direct RNA sequencing

RNA strands were either sequenced individually or pooled at equimolar amounts by length and encoding (Supplementary Table 1). Samples were prepared for direct RNA sequencing using the Oxford Nanopore Technologies (ONT) direct RNA sequencing kit (ONT, SQK-RNA002) and protocols provided by ONT (protocol version: direct-rna-sequencing-sqk-rna002-DRS 9080 v2 revO 14Aug2019-minion). Briefly, RT Adapter (ONT, SQK-RNA002) was ligated to 500 ng of each individual RNA sample or pool using T4 DNA ligase (NEB, M0202) for 15 minutes at room temperature. Reverse transcription was then performed with SuperScript III Reverse Transcriptase (Thermo Fisher Scientific, 18080044) by incubation at 50°C for 50 minutes, 70°C for 10 minutes, and then a 4°C hold. Reverse transcription reactions were cleaned up using 1.8x RNAClean XP beads (Beckman Coulter, A63987) and 70% ethanol. Reverse transcribed samples were eluted in nuclease-free water and quantified using Qubit 1x dsDNA high sensitivity assay (ThermoFisher Scientific, Q33230) or Qubit dsDNA high sensitivity assay (Thermo Fisher Scientific, Q32851). Next, reverse transcribed samples were ligated to the RNA Adapter (ONT, SQK-RNA002) using T4 DNA ligase (NEB, M0202) at room temperature for 15 minutes and then purified using 1x RNAClean XP beads and Wash Buffer (ONT, SQK-RNA002). Final libraries were eluted in 21 uL Elution Buffer (ONT, SQK-RNA002) and quantified using Qubit 1x dsDNA high sensitivity assay or Qubit dsDNA high sensitivity assay. Each library was sequencing on a R9.4.1 MinION flow cell (ONT, FLO-MIN106D) run on a GridION Mk1 sequencing device (ONT, GRD-MK1). Details about software, basecallers, and basecalling models used can be found in Supplementary Table 1.

#### A.2.4. Nanopore DNA sequencing

DNA strands were either sequenced individually or pooled at equimolar amounts by length and encoding (Supplementary Table 1). Samples were prepared for nanopore DNA sequencing using the LSK110 ligation sequencing kit (ONT, SQK-LSK110) and protocols provided by ONT (protocol version: genomic-dna-by-ligation-sqk-lsk110-GDE 9108 v110 revV 10Nov2020-minion).

Briefly, 200 fmol of individual DNA fragments or pools were mixed with 3 *µ*L NEBNext Ultra II End Prep Enzyme Mix and 3.5 *µ*L Reaction Buffer (NEB, E7180S) and incubated at 20°C for 5 minutes followed by 65°C for 5 minutes to repair fragment ends. DNA was then purified using 1x AMPure XP beads (Beckman Coulter, A63881) along with 70% ethanol, eluted in 61 *µ*L nuclease-free water at room temperature, and quantified with the Qubit 1x dsDNA high sensitivity assay. End-prepped DNA was then mixed with 25 *µ*L Ligation Buffer (ONT, SQK-LSK110), 10 *µ*L NEBNext Quick T4 DNA ligase (NEB, E7180S), and 5 *µ*L Adapter Mix F (ONT, SQK-LSK110). Reactions were incubated for 15 minutes at room temperature. DNA was purified using 1x AMPure XP beads and Short Fragment Buffer (ONT, SQK-LSK110). DNA was eluted in 15 *µ*L Elution Buffer (ONT, SQK-LSK110) at 37°C for 10 minutes. DNA was then quantified with the Qubit 1x dsDNA high sensitivity assay. Each library (50 fmol) was sequencing on a R9.4.1 MinION flow cell (ONT, FLO-MIN106D) run on a GridION Mk1 sequencing device (ONT, GRD-MK1). Details about software, basecallers, and basecalling models used can be found in Supplementary Table 1.

### A.3 Calculating Byte Error Rate

We calculate the byte error rate *P*_*l*,*s*_ for a given byte position *l* in an encoded strand *s* that with |*R*| sequencing read samples as:

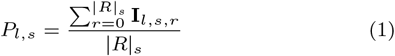

Where **I**_*l*,*s*,*r*_ is an indicator function that evaluates to 1 when there is a different byte compared to the original data in read *r* at byte position *l* for strand *s*, and 0 otherwise. To aggregate the byte error rates across encoded strands to determine general trends for a given design across byte positions, we take the arithmetic mean of Equation 1 across all strands (|*S*|) in a design as follows:

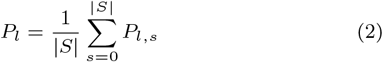

When considering a total aggregate byte error rate over all positions and encoded strands to determine a singular error rate that can be used to calculate outer code requirements we average Equation 1 over both the total number of positions |*L*| and the strands that have been encoded for a given design:

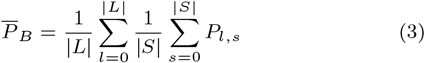

Equation 3 can be interpreted as an estimate of the byte error rate that would be generated if all bytes of all encoded strands and positions were equally likely to be chosen at random assuming that each position and strand are equally represented in the population. Thus, it represents the amount of information that can be recovered on average for each byte and indicates the information capacity of the channel for a given combination of inner code and decoder. This in turn can be used to determine additional redundancy for the inner code that is necessary to ensure a high reliability for recovering a data set of a certain number of bytes.

### A.4 Byte Error Rate Measurement Error

Given that we take a random sample of reads to generate a data set that can make computational times reasonable, our measurements of *P*_*l*,*s*_ are thus a random variable. The value of *P*_*l*,*s*_ is a sample of the mean of the Bernoulli random variable **I**_*l*,*s*_ that is 1 when an error occurs in strand *s* at position *l*. We assume that all *P*_*l*,*s*_ are normally distributed given our methodology for sampling strands. A general rule that allows the normal distribution assumption to be made for the proportion is that *np* ≥ 5 should be satisfied for *n* samples and a proportion *p*. Given the error rate is the order of 10% and number of samples for the baseline HEDGES decoding algorithm is 100k, the *np* ≥ 5 threshold is easily met. Beam Trellis algorithm error rates were shown to be on the order of 1% and thus 0.01 * 2000 = 20, and likewise across all strand designs studied for the Alignment Matrix algorithm error rates ranged on the order of magnitudes of [0.001 − 0.01] for a sample size of 10^4^ and thus also satisfies the threshold.

Thus, by linearity of expectation, both *P*_*l*_ and 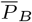 of Equations 2 and 3 are normally distributed with means that are averages of the true means of the **I**_*l*,*s*_ Bernoulli random variables. To calculate the variance of *P*_*l*,*s*_ we use ***Var*** (**I**_*l*,*s*_)*/*|*R*|_*s*_ which we estimate as follows:

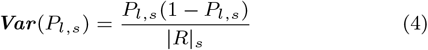

Then we can calculate the variance of *P*_*l*_ as:

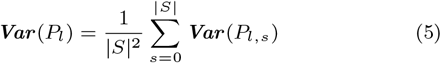

Here we assume that byte errors are independent across strands. In the case of 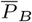 this is not the case as we have seen that byte error rates *within* a strand for the Alignment Matrix and *HEDGES* decoding algorithms are not independent of previous bytes. That is, if the error rate increases for *P*_*l*,*s*_ we expect *P*_*l*+1,*s*_ to increase as well. Therefore, we include covariance terms between byte positions in the calculation of 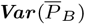 as follows:

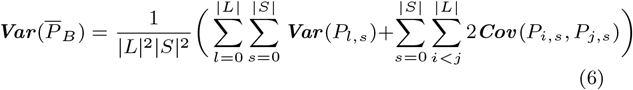

The value of ***Cov*** (*P*_*i*,*s*_, *P*_*j*,*s*_) is the covariance of sample means of the Bernoulli random variables of **I**_*i*,*s*_ and **I**_*j*,*s*_, which can be written as:

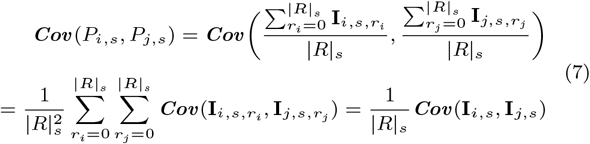

Here we have used the bilinear property of covariance to obtain an equation for the covariance of the measured byte error rates in terms of the covariance of the Bernoulli random variables which we can estimate using the following relationship between expected values and covariance:

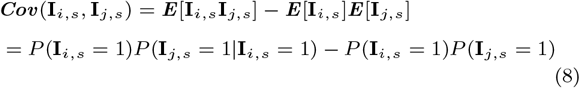

The value of *P* (**I**_*i*,*s*_ = 1) is the probability of a byte error rate at position *i* in strand *s*, which we can estimate with *P*_*i*,*s*_. The conditional probability *P* (**I**_*j*,*s*_ = 1|**I**_*i*,*s*_ = 1) captures the probability that a byte will be in error given the preceding byte registered an error (*j > i*). We do not measure this conditional probability directly, but a conservative estimate of the covariance and the following overall variance can be made by assuming *P* (**I**_*j*,*s*_ = 1|**I**_*i*,*s*_ = 1) = 1 which will result in a covariance that is larger than the true value. Finally, we arrive at our estimation of covariance ***Cov*** (**I**_*i*,*s*_, **I**_*j*,*s*_) = *P*_*i*,*s*_ − *P*_*i*,*s*_*P*_*j*,*s*_. With a calculation for the variances of each average error rate used, we use each mean ±1.96 standard deviations to calculate our error bars.

### A.5 Calculating Overall System Density

We determine the total density of a storage system as the product of four information rates as follows *ϕ*_*T*_ = 2*ϕ*_RS_*ϕ*_overhead_*ϕ*_HEDGES_. Here the factor of 2 represents the maximum capacity of information that can theoretically be stored per base, *ϕ*_RS_ represents the rate of the Reed-Solomon code used to withstand errors that persist after decoding the inner code, *ϕ*_overhead_ represents the ratio of information carrying bases to the total number of bases that need to be stored when implementing the strands needed for the storage system, and *ϕ*_HEDGES_ is the rate of the inner code used to translate digital information into bases that construct the molecules of the DNA storage system. While other terms require some calculation, the *ϕ*_HEDGES_ term is simply the rates that were chosen as parameters for this study.

The value of *ϕ*_*overhead*_ needs to consider strand length, the amount of that strand which will carry information corresponding to the raw information to be stored, and the different portions of the strand that will be needed to effectively recover information from the storage system. Implementing a DNA storage system typically requires an indexing region on each strand to allow for the information that is retrieved from unorganized sequencing data to be placed in the proper order. Furthermore, functional bases may be required within a strand to enable functionality such as PCR, transcription, etc. We calculate *ϕ*_*overhead*_ with the following formula for a molecule of *L* bases:

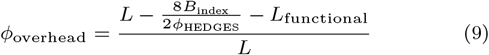

There are several parameters that need to be chosen in Equation 9. *B*_index_ represents the number of bytes that are used for indexing strands. In this work we choose *B*_index_ = 4bytes which allows for more than 4 billion molecules to represent some amount of data. Even for molecules of just 100 bp, 4 billion molecules surpasses the total base output of high performance flow cells such as PromethION Flow Cells. While this amount of indexed molecules may limit the amount of data that can be indexed together, it has been shown that physically addressing molecules through PCR (Organick et al. (2018)) and multi-stepped PCR (Tomek et al. (2019)) is a viable solution to increase the address space of a single pool of molecules by thousands (Organick et al. (2018)) to potentially millions (Tomek et al. (2019)) of additional levels of addressing. The bases associated with this addressing is modeled using the *L*_functional_ term which we set to 137 bp, the number of bp that are used as buffers/promoters in our strand designs. Given that the primers used in the aforementioned molecule addressing strategies are approximately 20 bp, *L*_functional_ = 137bp offers a sufficient budget for multiple primer addresses in the assumption that the buffer/promoter regions are simply repurposed.

Calculating *ϕ*_RS_ requires finding the minimum Reed-Solomon parity required to reliably recover a set of data. In this work we assume RS codes over GF(2^8^), and thus codewords of length 255 = |*D*| + |*P* | with |*D*| data symbols (bytes) and |*P* | parity symbols. Once a sufficient |*P* | is determined, the density of the RS outer code becomes 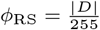 .

To come to a conclusion on what Reed Solomon code design should be chosen, we use 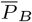 to calculate the probabilities of decoding success of a single codeword and then project that probability to a larger storage system. This is done with Equation 10. This equation sums up the probability of events that will cause a RS codeword to fail decoding, e.g. 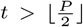 for *t* corruption errors in a RS codeword consisting of 255 = *D* + *P* total symbols with *P* redundant symbols and *D* data symbols. Considering each RS codeword must decode successfully, and assuming each is independent of each other, the mean time to failure (MTTF) will be 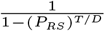.

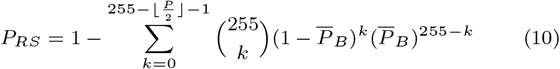

With each individual rate component known, the following equation can be used to combine them to determine an overall bits/base density of a storage system:

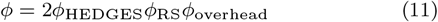

### A.6 CTC Models

For DNA reads, Beam Trellis and Alignment Matrix algorithms are integrated with Oxford Nanopore Technology’s open source basecaller Bonito (https://github.com/nanoporetech/bonito). Our implementations extend this code base starting from commit d36dfca. We use the CTC architecture provided with Bonito, with model version dna_r9.4.1@v2. For RNA reads, we use open source basecaller and model RODAN retrieved from (https://github.com/biodlab/RODAN), and our implementation is based on commit 029f7d5.

### A.7 Preprocessing FASTQ Reads

In our analysis, we post-process the raw sequencing reads from nanopore devices in order to identify which sequencing read belongs to each encoded strand, and to remove low quality reads that are unlikely to provide useful information regarding byte/base error rates. Our process is outline in Figure 2. We start with the FASTQ files of nanopore reads provided by the ONT basecallers, and pass these through FrameD’s sequencing read mapping utilities Volkel et al. (2023). The output of this step provides a map that links sequencing read IDs with the indexes of the encoded strands. We then take this mapping information and create a random subset of reads for each individually encoded strand. These subsets are used to partition the original FASTQ file and the raw electrical data that is stored in the FAST5 format to create the set of reads that are used for the decoding analysis that results in byte/base error profiles.

During the initial analysis of the ONT basecall data to create the index/read ID map, we pass each read through a series of filters. The first filter we apply is an alignment filter between the sequencing read and the 3^′^*/*5^′^ buffer regions that we apply to the ends of each encoded strand. The buffer regions are intended to provide protection from skipped bases that may occur on the transient entrance/exit of a molecule traversing the nanopore, but we also require their removal prior to decoding by the HEDGES algorithm. This is because HEDGES will not recognize the additional buffer regions as part of the overall message, and so decoding will likely fail. In order to ensure that the alignment is a reasonable quality, we require that each alignment has at least 70% matching bases within the alignment. Following the alignment, the region of the sequencing read that is aligned to is removed and the subsequent remaining read’s length is inspected. If the read is either too long, or too short, we remove it from consideration with the reasoning that extremely short reads indicate incomplete reads that are highly unlikely to decode and long reads indicate chimeric read events. We avoid including these short or long reads with the goal of avoiding anomalous reads in the inclusion of base/byte error analysis so that typical nanopore behaviour that results in decoding failures can be reasoned about. For the lower bound, we use a value that is 6-7% shorter relative to the length of the region that encodes information, and for the upper bound we use a value of 300-400 bases longer relative to the length of the region of the molecule that encodes information. As the final step, we use the HEDGES algorithm to decode the index of the read, and as a final check to ensure that the index bytes are decoded correctly we calculate a CRC check byte to avoid incorrect indexes being attributed to sequencing reads.

**Fig. 2:**
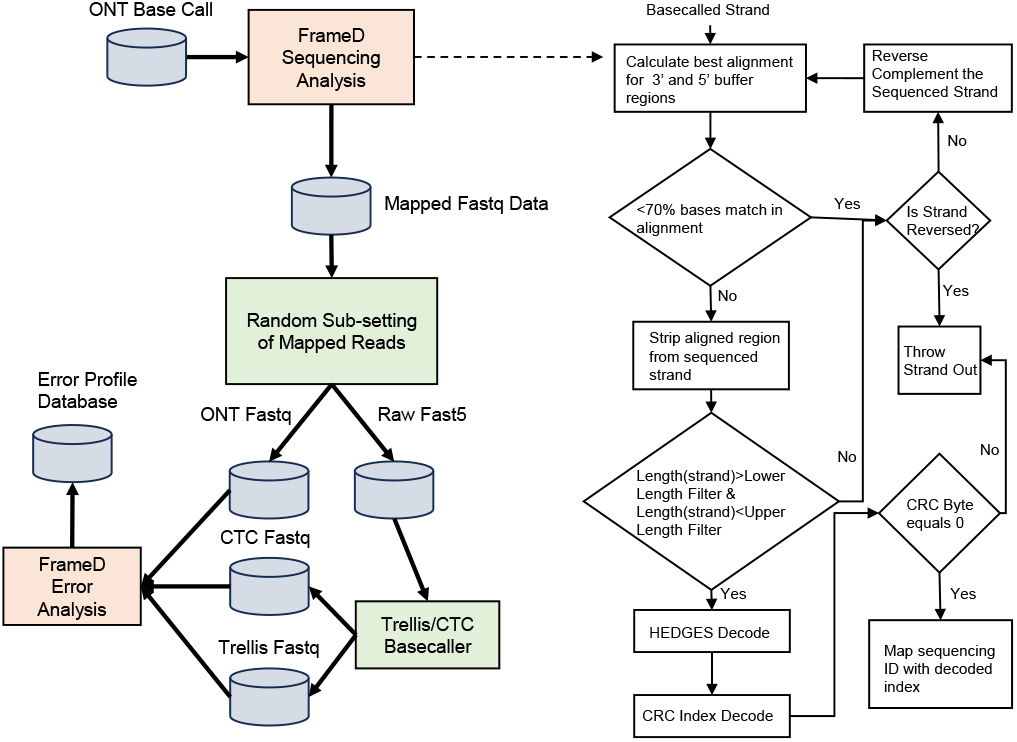
Diagram of our workflow for determining a working set of nanopore reads to evaluate against our decoding algorithms.

#### A.7.1 Basecalling Error Rates

As supporting evidence for our sequencing post-processing methodology, we calculate the edit-rate for each strand of the 0.167 HEDGES rate as shown in Figure 3. We calculate the edit rate by determining the edit operations used to transform the originally encoded strand to the read observed during sequencing after removing buffers (e.g. only HEDGES encoded bases). By not including buffers, we are able to directly see what error rates the error correction code is being faced with. We then accumulate the number of edits and type at each base position to determine a base error rate. During analysis we found that short reads manifest with high numbers of deletion errors towards the terminal base positions of the encoded strand. To account for noise in the edit rates, we average over a rolling window of 50 bp to allow for local variation and global trends across reads to be observed. From this data we can concluded that deletion error rates (and all other rates as well) remain consistent across the strand. This indicates that there is not a large positional influence in error rate across strand positions due to sequenced molecule length.

#### A.7.2 Q Score of Partitioned Reads

To further investigate the impact of our filtering on the quality of reads kept for analysis we study the impact on the distribution of quality scores determined by the ONT basecallers. Figure 5 shows quality score distributions for reads covering all density and length design points covered in this work. The line annotated as *Total* represents the quality score distribution for every read in the sequencing run, other lines are labeled with encoded strand identifiers and these distributions represent the distributions for each read *after* they have passed through the filtering steps of Figure 2. We also label the plots with two means of each distribution. The first mean is the mean of the entire distribution, and the second mean represents the mean of the distribution if all Q scores less than 9 are ignored. The cutoff point of 9 was chosen based on ONT’s default setting of determining poor quality reads. Based on the average of the entire distribution, the filtered reads generally have an average that is larger than the total population. However, it is apparent that this shift is a result of a large number low quality reads between Q score of 5 and 6 that do not exist in the filtered read sets. If we consider a Q score cutoff of 9, we can see that the averages of the total distribution and the filtered sets are considerably closer. Furthermore, we can also observe that the peaks of the filtered distributions do not bias towards the high Q score tail of the total distribution, indicating that our filtering process does not have a preference for very high quality reads. Provided that our filtering keeps reads close to the desired read length, we can infer that the low quality score reads lost in our filtering correspond to short-length reads and further motivates eliminating short reads from consideration.

**Fig. 3:**
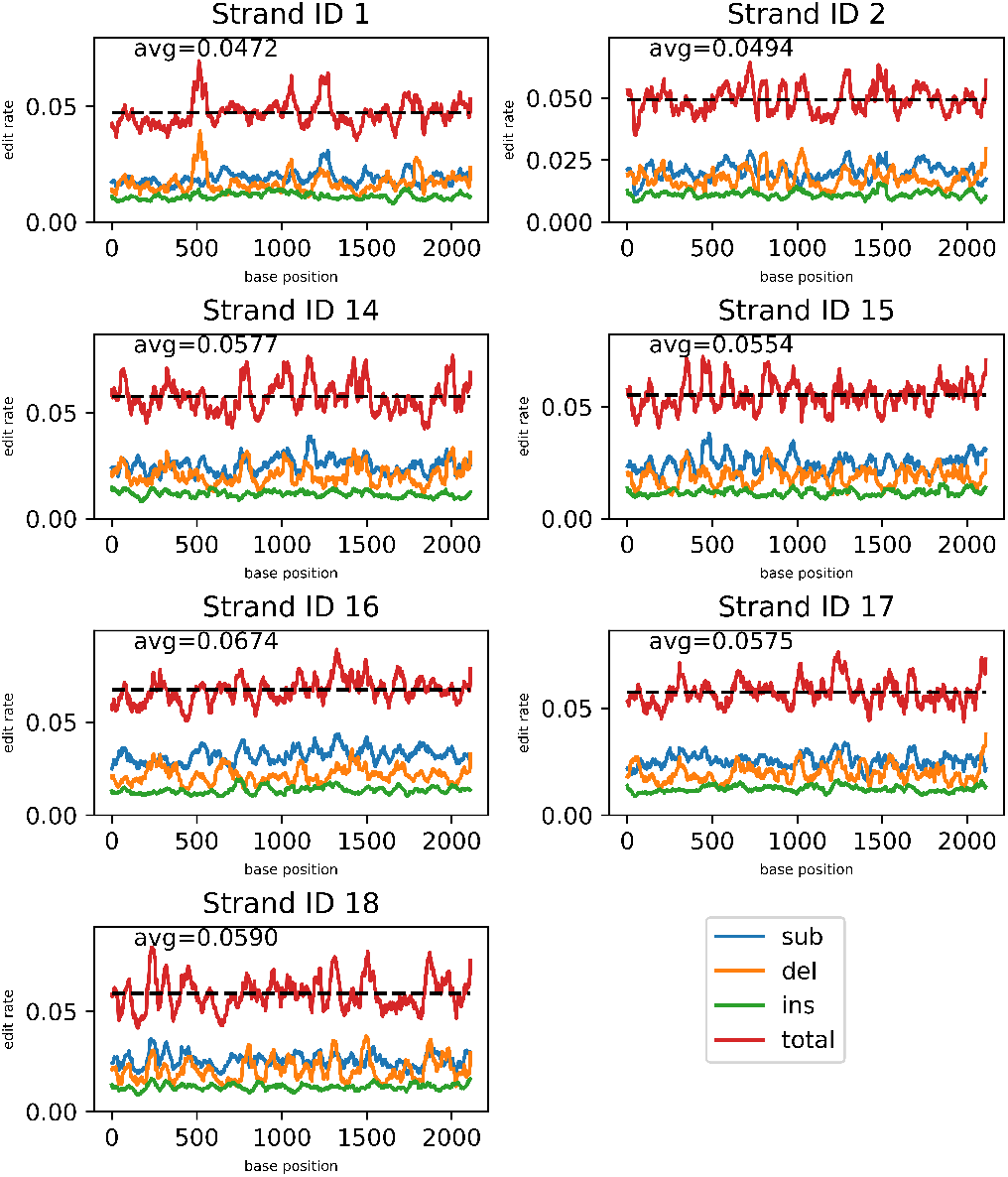
Basecalling error rates measured for each strand of the 0.167 HEDGES rate design of length 2297 bp over a sample of 100k reads for each strand. Error rates are measured over the 2160 bp that are carrying the encoded information. Each line represents a rolling arithmetic mean for errors over 50 bp windows for substitutions (sub), deletions (del), insertions (ins), and the total number of errors. Dashed horizontal line represents the average error rate over the entire strand calculated by summing all errors at all positions and dividing by the total number of reads and positions.

There was some cross-batch variation in quality scores. For example, the average quality score for 0.5 rate HEDGES was significantly lower (10.32) than that of the 0.125 rate (13.13). While the encoding densities are different, the constraints imposed on the encoder are the same, e.g. we only allow for homopolymers of length 3 at most, a fixed GC content to be 50%, encoded length ≈ 2160. Also, we observe that there is no significant change in Q score across the 5 different strands of the *HEDGES 1667 Length full* run. Thus, we attribute the quality score variation to variations in the nanopore flow cells.

**Fig. 4:**
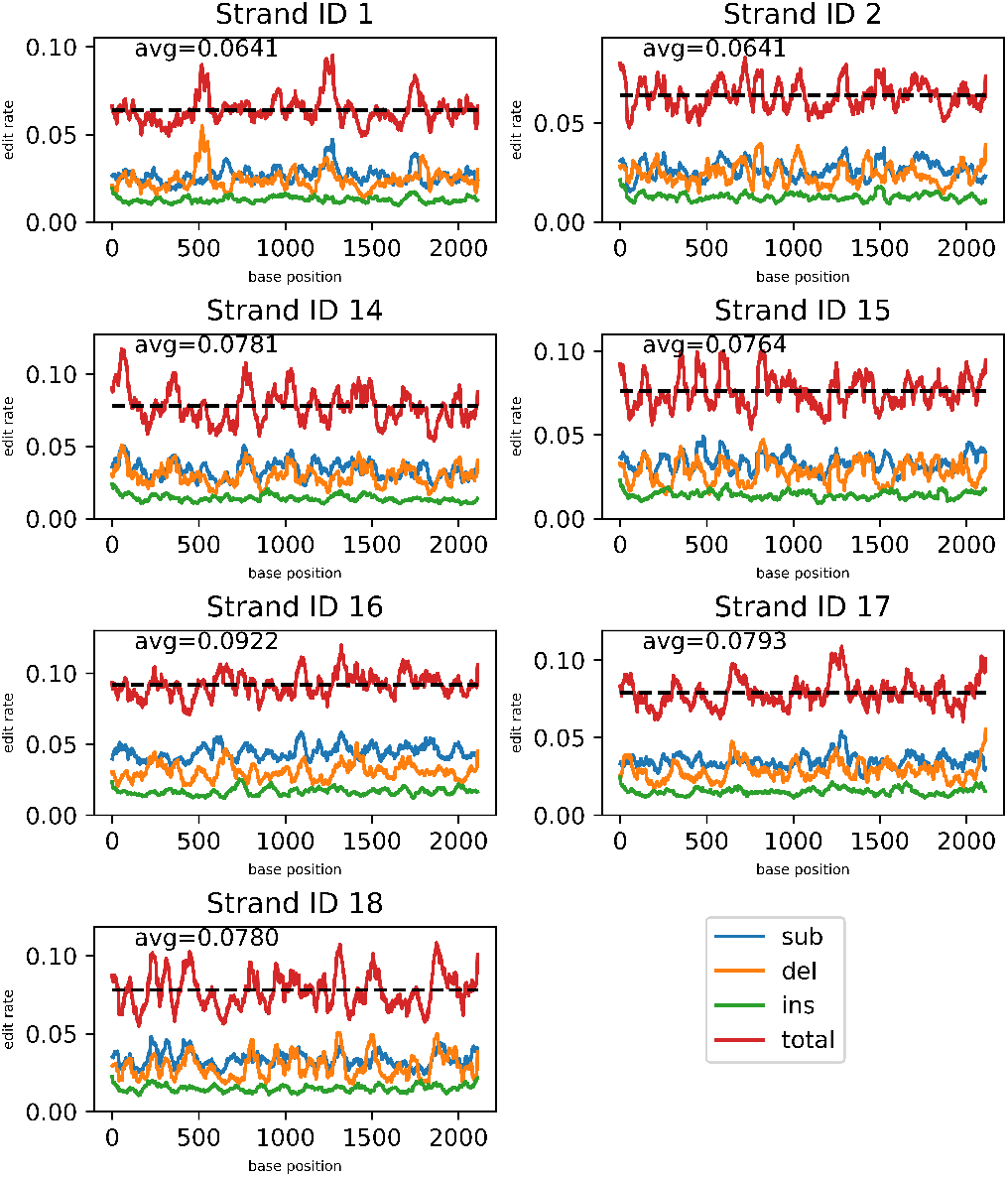
Rolling average base edit rates over a 50 bp window for the open source Bonito-CTC basecaller applied to the 0.167 HEDGES library across 100k reads.

### A.8 Preprocessing CTC Matrices

To utilize the decoding algorithms that leverage CTC scores for decoding HEDGES codes, there are several steps that need to be completed beforehand. Figure 6 illustrates the preprocessing required to transform electrical data into the proper CTC form for our studied algorithms. First, the FAST5 data needs to pass through a CTC-based machine learning (ML) model. In the case of DNA molecules, this model is an open source CTC version of the Bonito basecaller, for RNA strands the model is the open source RODAN ML model Neumann et al. (2022). These models generate the CTC matrices that are required to decode each nanopore read.

As mentioned, there are bases which serve molecular function purposes that surround the payload that we are interested in studying the decode error rates of. Thus, the CTC matrices provided by the ML models will also include scores related to these bases that are unrelated to the encoded payload. The CTC scores that correspond to the auxiliary bases are pruned so that the algorithms can make higher quality alignments for messages without noise introduced from alignments to non-data regions. Supplemental Figure 6 describes our workflow for pruning CTC information and understanding the read direction that the CTC information represents. The CTC pruning process performs alignment in both directions, forwards and reverse complement. This is especially important for DNA reads as any read can be forward or reverse complement, and we need to know which bases to generate in to properly decode relative to the CTC data. However, for RNA we only need to consider alignment in the forward direction since it is only a single strand molecule. Once alignment is complete, if the reverse score was found to be larger we reverse the direction of the CTC matrix and set the HEDGES base-generation to emit complements. We finally are now able to perform trellis decoding for the HEDGES strands using CTC scores during which we generate a FASTQ basecall file with the final decoded message.

**Fig. 5:**
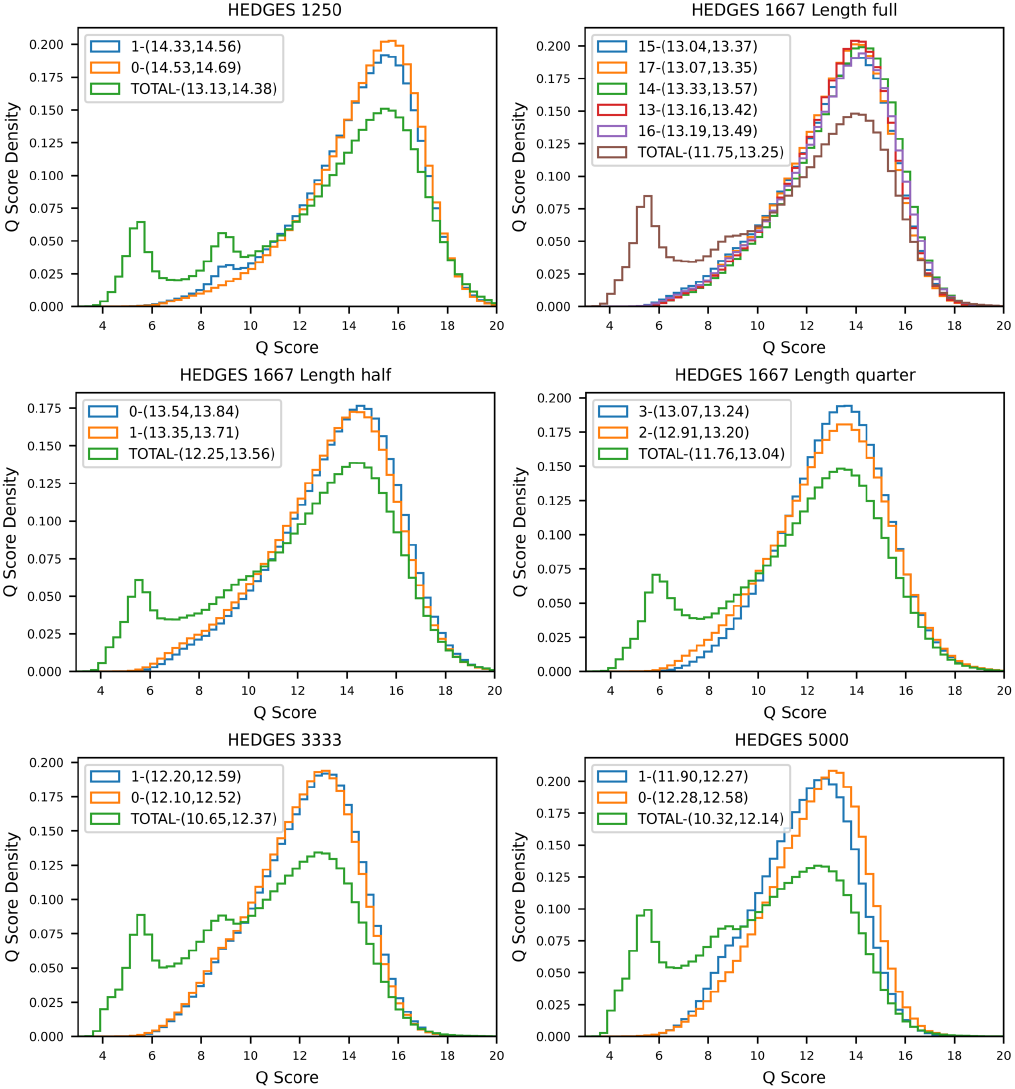
Q score distribution for filtered and all reads for sequencing runs performed for each strand design.

**Fig. 6:**
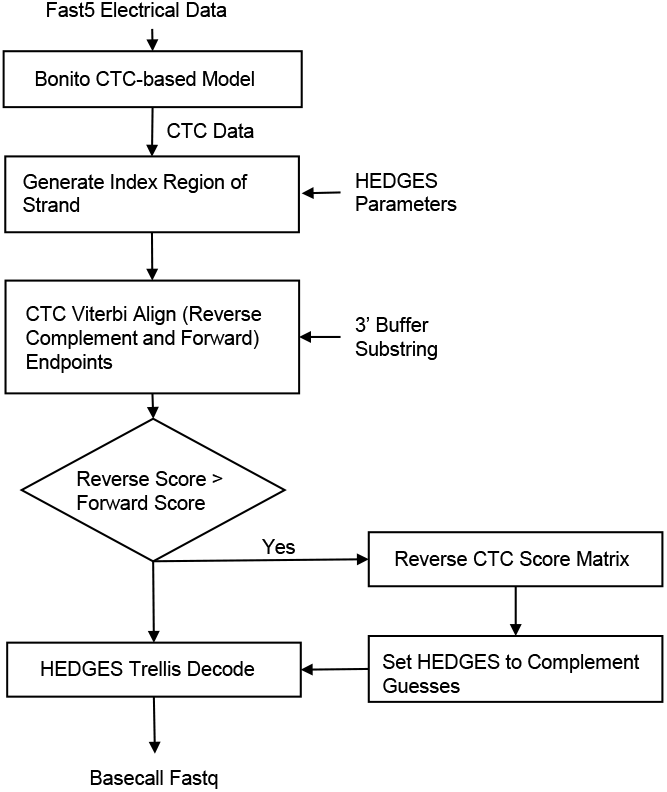
Processing flow of electrical data stored in FAST5 files throughout the process of decoding HEDGES encoded strands using both the Alignment Matrix and Beam Trellis decoders.

The process of aligning a known base sequence to a CTC matrix is a common strategy referred to as *forced alignment* that has been applied in speech recognition to determine regions of the CTC scores that correspond to utterances. In our case, the utterances will be the buffer/index regions of our DNA/RNA strands, and so we follow a procedure similar to that provided by Kürzinger et al. (2020), which is outlined in Figure 7. The algorithm is set up as a trellis, with each state corresponding to a base in the known sequence, or a blank that is inserted between each base. An initial state *ϵ* is included to allow for alignments to be found at any starting time point in the CTC matrix. The trellis states are evaluated for each time step in the CTC matrix, with edges between states that correspond to valid CTC encodings of the know base sequence. For example, if the known base sequence has a repeat such as *GG* then there cannot be a direct edge between each of the *G* states in the trellis. There must be blank symbol in between. This is not necessary for non-repeats as shown in Figure 7 with the edges connecting *A* to *G*.

**Fig. 7:**
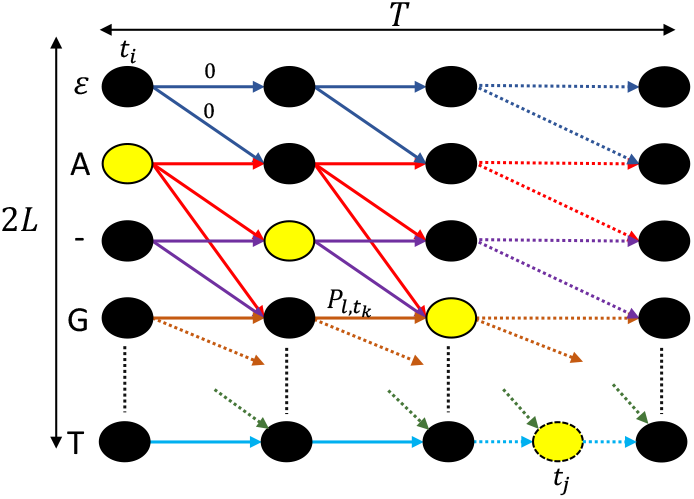
Trellis used in the process of aligning buffer regions using CTC scores for the purpose of determining start points for the HEDGES decoders.

The score of each state is calculated as the maximum of the score of all incoming states summed with the log-probability of the state’s symbol at the given time point. The following equation provides the score *P*_*l*,*t*_ for state *l* at time *t*.

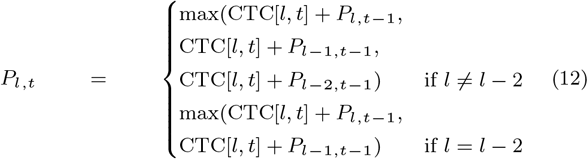

The first case of Equation 12 is for the case of non-repeats between two consecutive bases in the sequence, and the second is for the repeat case. With this definition for scoring, the trellis can be filled out so that each state is assigned a back-trace path corresponding to the edge taking during the maximum of Equation 12. After this is done, the starting point for the back trace pass is taken as *t*_*j*_ = arg max_*i*_(*P*_2*L*−1,*i*_), the time step that results in the maximum score for the final base in the known sequence. Starting from here we back trace until the first base in the sequence is reached, e.g. following the highlighted states until *t*_*i*_ in Figure 12. Thus, the time steps from *t*_*i*_ to *t*_*j*_ are taken as our alignment.

### A.9 GPU Implementations

#### A.9.1 GPU Beam Trellis Port

We provide a general outline here of our GPU port of the work by Chandak *et al*. Given that GPU implementations of algorithms are highly dependent on several design choices such as thread layout, memory layout, and resource usage such as local shared memory spaces, we aim to show that the choices that were made for the GPU implementation of the Beam Trellis algorithm are reasonable and should not hinder the algorithms performance relative to our Alignment Matrix algorithm. An outline of our design choices is presented in Figure 8. Our implementations are based on the NVIDIA CUDA GPU programming platform.

**Fig. 8:**
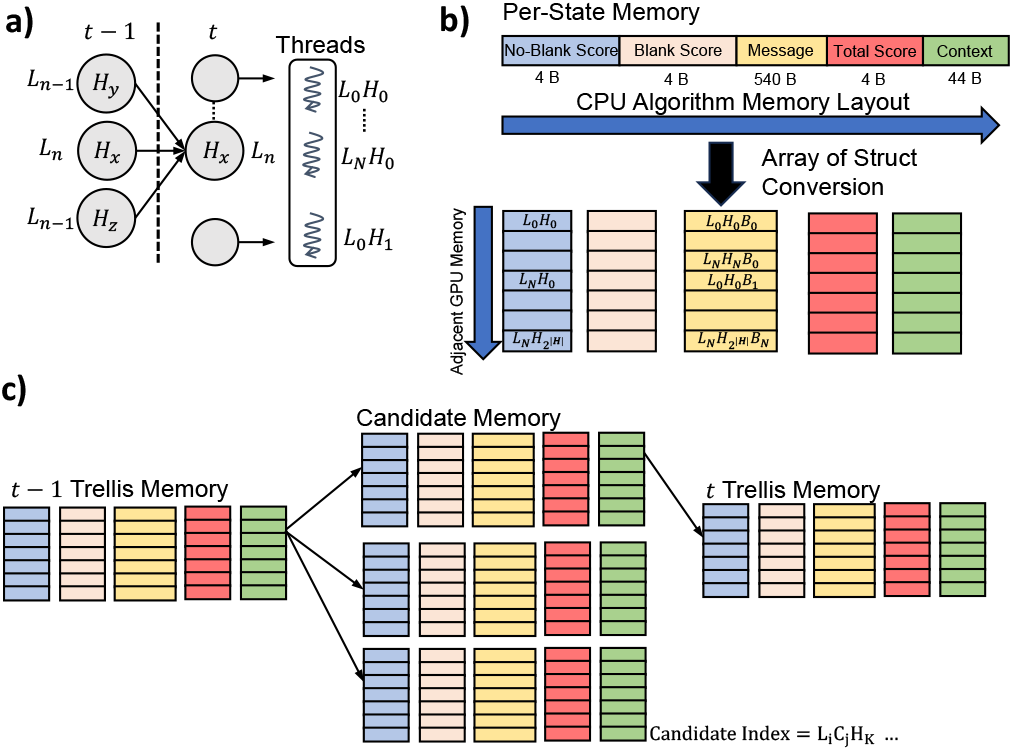
**a)** Trellis to thread mapping used for constructing parallel work units for the Beam Trellis algorithm. **b)** Memory layout of data related to each state at trellis time step *t* − 1. **c)** Layout of temporary candidate data that is used to determine the best incoming path to choose for a given state.

The first step we take in constructing our GPU implementation for the Beam Trellis algorithm is choosing the parallel units of work and their layout on the GPU device. For trellis-based algorithms, the computation for each state’s score at a given step are all independent of each other. Also, given that there are a large number of states for the Beam Trellis algorithm, ≈ 2000 × 256 = 512000, parallelization over the states will easily keep a GPU device busy. Thus, as shown in Figure 8 we assign a thread to each state, and we order each thread co-lexicographically over the number of length states *L* and convolutional states *H*. That is, for a strand length of *N* and number of convolutional states *M*, thread indexes are ordered according to *L*_0_*H*_0_, *L*_1_*H*_0_, …, *L*_0_*H*_1_, …*L*_*N*−1_*H*_*M*−1_. This ordering becomes important when considering memory access patterns of threads and maximizing the amount of read coalescing that can impact memory throughput of the implementation.

The second step for our GPU implementation was to reconfigure the memory layout of data structures to enable GPU memory access coalescing. The core data structure that we associate with each state is shown in Figure 8b. This includes 5 different fields: ***No-Blank Score*** a 4-byte float that stores the aggregated scores of all paths to a given state that do not end in a blank, ***Blank Score*** a 4-byte float that stores the aggregated scores of all paths to a given state that do end in a blank, ***Message*** a 540-byte bit-array that stores the base-message using 2 bits per base that corresponds to a given state for a given trellis-step, ***Total Score*** a 4-byte float aggregating non-blank and blank scores, and ***Context*** a 44-byte object that is responsible for keeping track of which base to emit based on tracked constraints such as GC balance and homo-polymer runs. Typically, for a CPU implementation, these fields should be adjacent when evaluating each state because there will be high temporal locality between the fields. However, because GPU threads all execute identical instructions in lock-step given their single instruction multiple thread (SIMT) execution model, a GPU device’s memory throughput is better optimized when each thread accesses adjacent memory locations. Thus, similar to how we order threads, memory locations are co-lexicographically ordered over *L*_*i*_*H*_*j*_. Note, because the message field is so large (540-bytes) and thus will contribute to a large portion of the memory bandwidth, we further re-arrange the bit-array to be co-lexicographically ordered over *L*_*i*_*H*_*j*_ *B*_*k*_ where *B*_*k*_ indicates a bit position in the bit vector.

To see why arranging threads and memory in this manner results in adjacent threads accessing adjacent memory we can analyze the memory access patterns for each thread with this given memory layout. We co-locate threads that have a common convolutional state *H*_*x*_. For example, *L*_0_*H*_*x*_, *L*_1_*H*_*x*_, …, *L*_*N*−1_*H*_*x*_ are adjacent. Each thread in this set will need to access the memory corresponding to three separate candidates, each corresponding to three different valid previous convolutional states (Figure 8a). Given that *H*_*x*_ is constant amongst these threads, they will all have the same incoming previous states. Furthermore, for some thread *L*_*n*_*H*_*x*_ in this set, the incoming candidates will have position portions of either *L*_*n*−1_ or *L*_*n*_. For *L*_*n*−1_ this means that the memory access according to each thread will be ∅, *L*_0_*H*_*y*_, …*L*_*N*−1_*H*_*y*_, which is a favorable access pattern for coalescing given the memory layout. The access ∅ is used for the case that there is no *L*_*n*−1_ candidate for *L*_0_. For *L*_*n*_, the state *H*_*x*_ does not change, nor the position, thus the access pattern becomes simply *L*_0_*H*_*x*_, *L*_1_*H*_*x*_, …, *L*_*N*−1_*H*_*x*_.

The memory accesses covered so far only concern with reading memory associated with incoming states of the previous trellis time step. Another memory region of concern is the memory used to store candidate results (Figure 8c). This memory holds the same memory fields as shown in Figure 8 b), but three copies are needed for each thread to account for each of the three possible incoming candidates. For this candidate memory, we lay it out such that the co-lexicographic order is based on *L*_*i*_*C*_*j*_ *H*_*k*_ This enables memory accesses to be coalesced since each state will fill in their candidates in the same order as every other adjacent state. That is, candidate coordinates of *L*_0_*C*_0_*H*_*x*_ and *L*_1_*C*_0_*H*_*x*_ will be written to at the same time during SIMT execution. We could also change the coordinates of candidate memory to be *L*_*i*_*H*_*j*_ *C*_*k*_, but because our implementation fixes each block and thus warp to have a constant *H*_*j*_ this reordering would be inconsequential to memory coalescing.

#### A.9.2 Alignment Matrix

The outline of the key optimizations done for the Alignment Matrix implementation is shown in Figure 9. First, we noticed that for the HEDGES algorithm there is a pattern where the transitions between states are self-transitions when the bits conveyed by a base are 0. That is, the state does not change when no new information is transmitted by a base. For this reason, we do not need to calculate edge scores as it is trivial to determine the best transition coming into a state for trellis steps *l* + 1 and *l* + 2 as indicated by Figure 9. Once a transition is reached between two trellis steps that require score comparisons to determine the maximum path (*l* + 2 and *l* + 3), we calculate the alignment score for the newly added base along with all bases accumulated along the 0 bit pattern paths. For example, the new base added on the transition between state *H*_*x*_ at step *l* + 2 going to *H*_*z*_ at step *l* + 3 is *T*, but the actual bases utilized in the Alignment Matrix calculation is *AGT* .

**Fig. 9:**
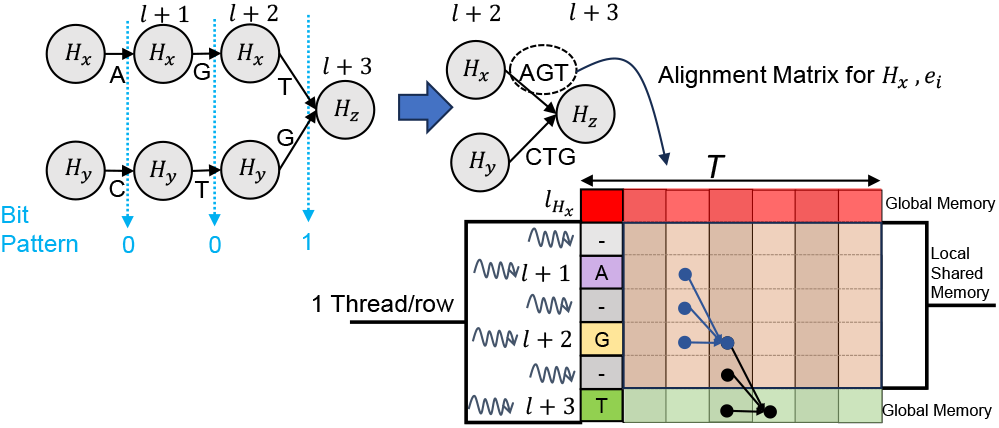
Outline of GPU implementation for the Alignment Matrix algorithm.

With the bases known on each transition, we can then calculate a score to compare two incoming edges reaching a state. This results in each edge requiring a calculation of its Alignment Matrix which is formed as shown in Figure 9. The top row of the matrix is a length *T* array of scores which corresponds to the alignment scores of the best path for state *H*_*x*_ at step *l*. These values are stored in global memory since they need to be utilized by succeeding steps in the trellis. Likewise, the final row for the last base needs to be stored in global memory since this row needs to be propagated along the trellis to facilitate future edge scores. However, there are a number of alignment scores that can be stored in the GPU’s shared memory since they are intermediate values that are not necessary for future calculations. These are the rows of the Alignment Matrix that correspond to bases that are not the last base in the edge’s complete string, or rows that correspond to blanks represented by *-* in Figure 9. This reduces the amount of global memory bandwidth, and also ensures that the required memory from the GPU is only proportional to *T* and not *L* × *T* which would be necessary if the entire Alignment Matrix was naively propagated through the trellis. Thus, the matrix alignment algorithm can maintain memory linear in strand length instead of quadratic since *T* = *O*(*L*).

Data flows in the Alignment Matrix from the first column to the last column, with each column corresponding to a time step which consumes the previous time step’s data as shown by the connection of matrix elements in Figure 9. Unfortunately, that means that calculating elements in successive columns needs to be serialized. However, this pattern also means that all rows in the same column can be calculated independently. This leads us to assign threads to each row for a given Alignment Matrix calculation. Thus, the number of threads launched per trellis step is *H* × *E* × *L*^′^ where *E* is the number of edges on a transition. Typically, *E* = 2 when 1 bit is transmitted per base, but provided that a base can represent up to 2 bits *E* = 4 is also possible for different HEDGES code rates which is supported by our implementation. *L*^′^ here is the number of independent rows that are being calculated for a transition edge. This depends on the HEDGES rate as well since lower code rates will increase the number of 0’s in the bit pattern which increases the number of rows calculated at each step. For a fixed strand length, *L*^′^ will impact run time such that lower density codes will decode faster than more dense ones. The reason is that more dense codes will not be able to leverage shared memory as much and require more global memory transfers.

### A.10 Soft Decoder Memory Comparisons

Based on both of our soft decoder implementations, we outline the calculations made to determine their memory consumption. Starting with the Beam Trellis algorithm, we calculate the total memory required with the following equation, assuming *N* for the total number of positions and *M* total convolutional states.

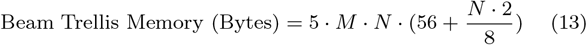

As we showed in Figure 8 b), each state requires 56 bytes plus the memory required for the message. Because the 66 bytes is constant with message size, we represent the message memory separately in Equation 13 as 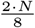 where we use 2 bits per base of a message. This quantity is multiplied by *M* · *N* because every state needs its own version, and another factor of 5 is included to account for 3 candidate copies and 2 copies required to store trellis time step *t* − 1 and time step *t* (Figure 8 c)). Using this equation we are able to project an estimate for memory as the number of states changes or the length of the strand changes.

The memory of the Alignment Matrix algorithm is allocated over a small set of critical arrays that are required to track alignment information, paths through the trellis, and context required by the HEDGES algorithm. The arrays we use are: a ***Context*** array of dimension 2 × *M* 44-byte data structure used to manage the HEDGES state between steps in the trellis, a *T* × *M* array of 4-byte floats named the ***Forward*** array that holds the alignment results across the all *T* steps of the CTC matrix for each of the *M* convolutional states (first row of the array highlighted red in Figure 9), a *T* ×*M* ×2 array of 4-byte floats named the ***Forward-Edges*** that tracks the alignment for each edge during a trellis transition (last row of the array highlighted green in Figure 9), and we utilize two *M* × *N* arrays of 8-byte values where one serves as the back-trace array after the trellis is complete and the other stores the bases that are emitted along the best-path edges. This leads to the following summation of memory for the Alignment Matrix algorithm as a function of the number of convolutional states and strand length.

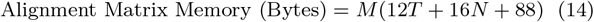

From Equations 13 and 14, it is clear that the Beam Trellis algorithm’s memory consumption is quadratic in strand length, while the memory consumption of the Alignment Matrix algorithm is only linear in strand length. Note, *T*, the dimension of the CTC matrix is generally not constant for each read even for the same strand length *L*. Thus, we require some assumptions for *T* in order to calculate the bytes required for the Alignment Matrix algorithm. We choose *T* to be the average CTC time dimension measured over the 400 reads used for benchmarking, which we measure to be 7400 as indicated in Figure 11 which displays the distribution of *T* values over these 400 reads. Given this assumption, we project to different strand lengths by scaling the *T* and *N* terms of Equations 13 and 14 by a factor of 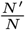 for some new strand length of *N* ^′^ bases. For our memory calculations, we fix *N* = 2160 bases and *M* = 256 convolutional states. Results for projecting memory consumption at different strand lengths is shown in Figure 10.

**Fig. 10:**
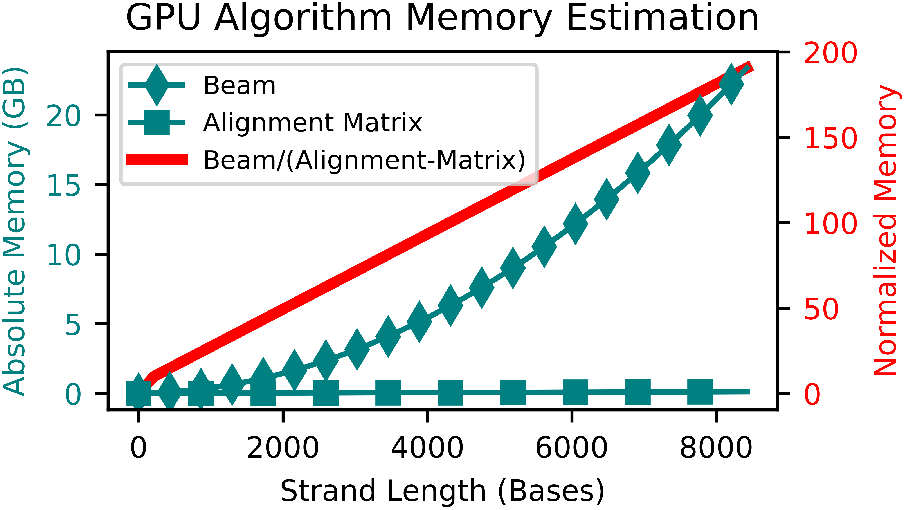
Average memory consumption comparisons between the Beam Trellis and Alignment Matrix algorithms.

**Fig. 11:**
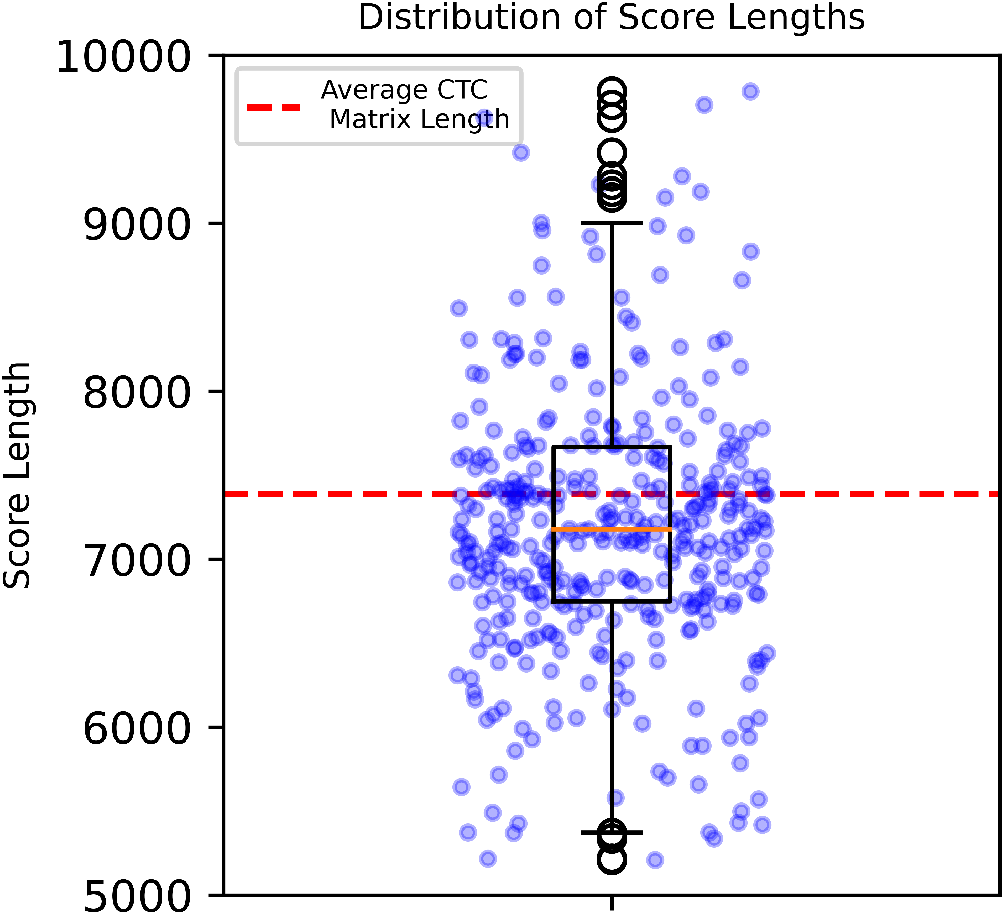
Distribution of CTC matrix time lengths for 400 reads that were analyzed in detail for run time and memory benchmarking.

## B. Additional Hard Decoding Analysis

Figure 12 shows the relationship between byte error rate versus position for each encoded strand when the ONT basecallers are decoded with the HEDGES hard decoding algorithm. This data confirms the behaviour observed by Press et al. (2020) of the Tree decoding algorithm where as position increases along the length of the strand the probability that the byte can be decoded decreases. The reason this occurs is due to running out of guesses before the read can be completely decoded. Increasing the number of guesses can decrease the byte error rate at each position, but the correlation between position and error rate still remains. We also note here that there is generally not a significant difference between each encoded strand’s byte error rate curves, or locations within each curve that have sharp changes in byte error rate. This indicates that events from the ONT basecaller that causes HEDGES to fail are not strongly correlated to any given position.

**Fig. 12:**
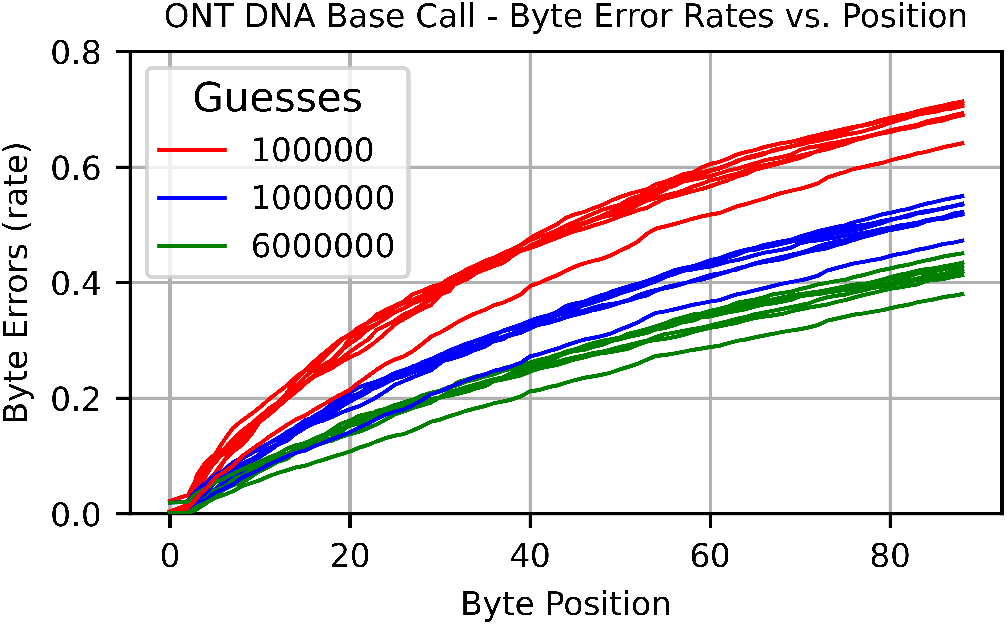
Byte error rate as a function of byte position when applying the HEDGES Tree search algorithm to ONT basecaller outputs for 3 different guess budgets. All curves correspond to the 1*/*6 encoding rate strands.

To show how byte error rates impact the cost for a given amount of reliability and storage system of a certain size, we plot a density heat map for various byte error rates and system sizes in Figure 13. In this analysis we assume a MTTF = 10^6^ in all cases. Based on this data, we show that the byte error rates associated with using the HEDGES tree based decoder result in no solution, or very low densities, when storage systems are larger than 1 TB. A scenario in which no solution occurs means that we exhausted all Reed-Solomon code designs in terms of the amount of the codeword allocated to data and parity symbols before the MTTF threshold was surpassed. Furthermore, even for small storage systems of 1 GB, densities are *<* 0.5 bits/bp for this decoder. These low densities confirm that the baseline hard decoder is not suitable for a single-read storage system in a nanopore sequencing setting.

**Fig. 13:**
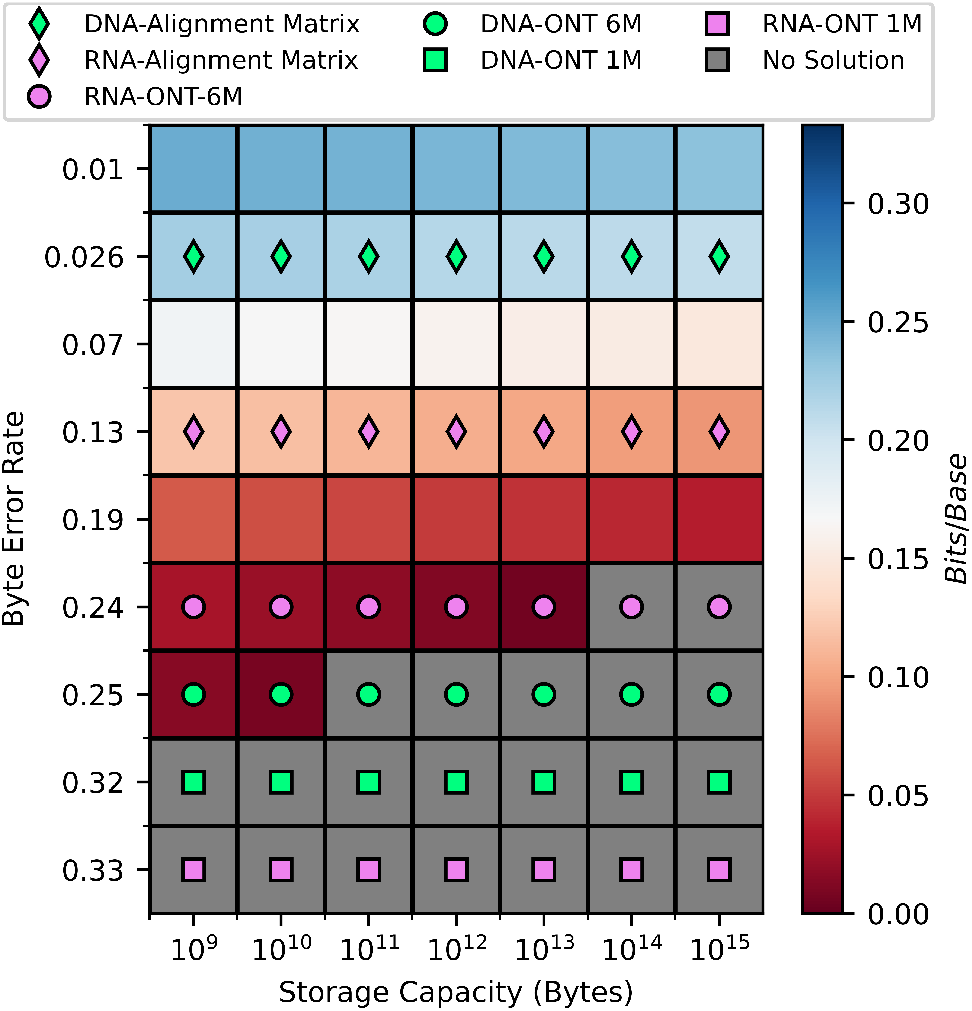
Byte error rate (rows) and storage capacity (columns) versus the achievable density (color bar) when assuming the 1*/*6 HEDGES rate code. Rows labeled with a marker indicate a byte error rate that was measured for that decoder. All DNA markers pertain to error rates measured on the full 7 strand library for the 1*/*6 rate HEDGES encoding. RNA/DNA-ONT *X*M refers to decoding RNA/DNA basecalls using *X* million guesses.

## C. Strand Length Analysis

Given that the shorter strands have been shown to have lower byte error rates, we are interested in understanding the error rate relationship as a function of byte position so that we may develop a method to project how strand length may impact other HEDGES densities. Figures 14 and 15 show how the byte error rate trends in relation to position when averaging over each individual strand and when considering strands individually respectively. Figure 15 shows that the shorter length strands have byte error rates that are similar on average to that of the full length strand at each respective position. We do note that the average byte error rate curves for the half and quarter length strands lie above and below that of the full length strand respectively. However, given that we have normalized the samples to contain reads between values of 15.1-15.4 of Q scores for both figures to factor out flow cell variation, we consider the deviation of the shorter strand byte error rates to be a product of how the model and decoder respond to different base sequences. Supporting this idea is Figure 14 which shows that there are full length strands that have individual byte error rate profiles that are close or that their error bars encompass the byte error rate of the individual short strands. We conclude from this data that an estimation of an error rate profile for the other studied HEDGES rates at shorter strand lengths can be derived by truncating their respective byte error rate curves appropriately.

**Fig. 14:**
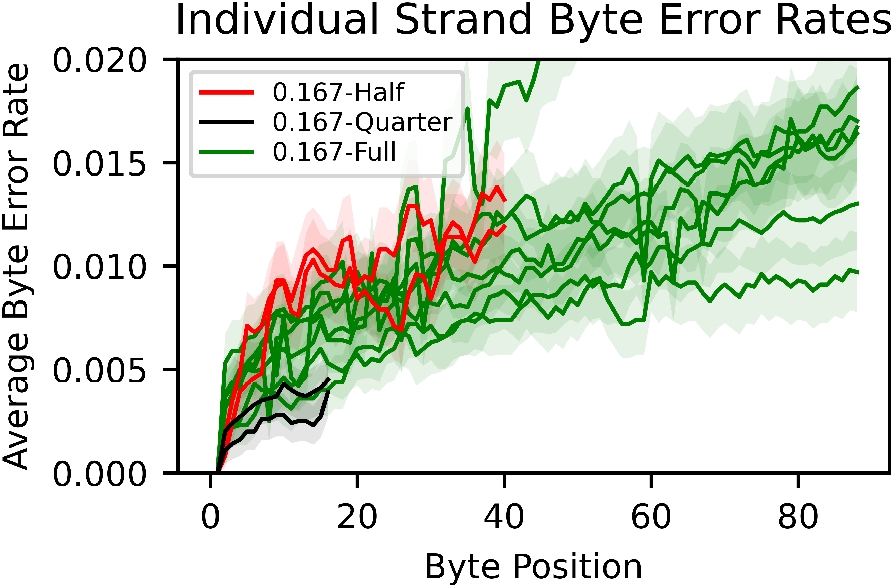
Byte error rate vs position for each strand across the three different synthesized lengths for the 1*/*6 encoding rate. Half, Quarter, and Full each refer to the three different strand length designs in increasing order. Error bars that represent a 95% confidence interval are calculated for each byte based on the variance calculation of Supplemental Equation 4.

**Fig. 15:**
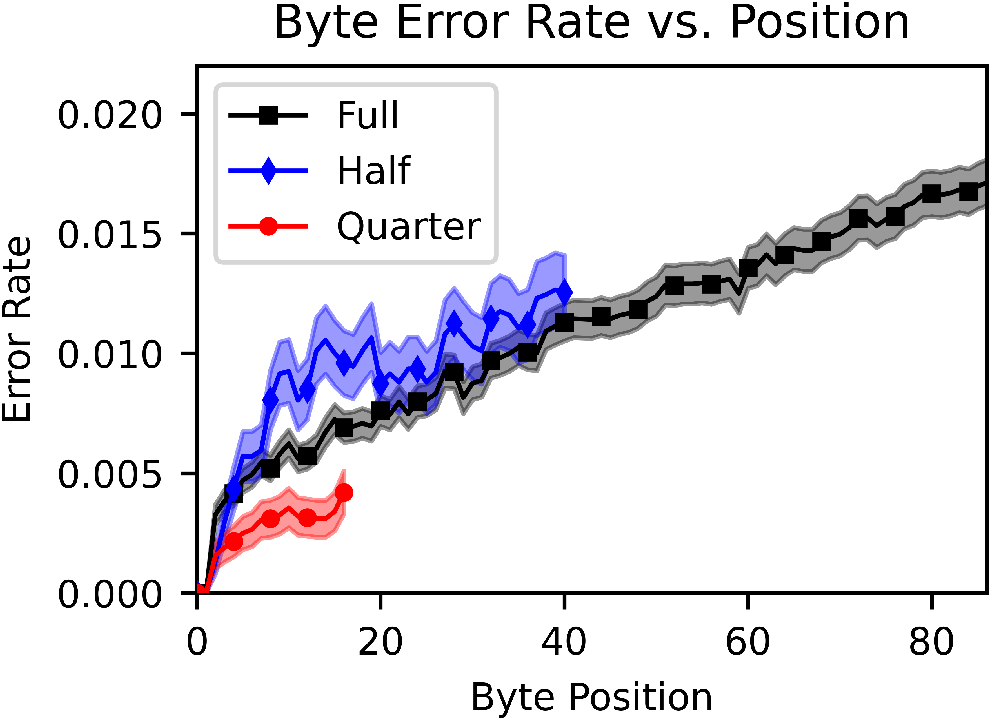
Byte error rate for each byte position average across all synthesized DNA strands for the 3 different lengths studied for the 1*/*6 rate HEDGES encoding. Error bars are calculated according to Equation 5 to represent a 95% confidence interval.

**Fig. 16:**
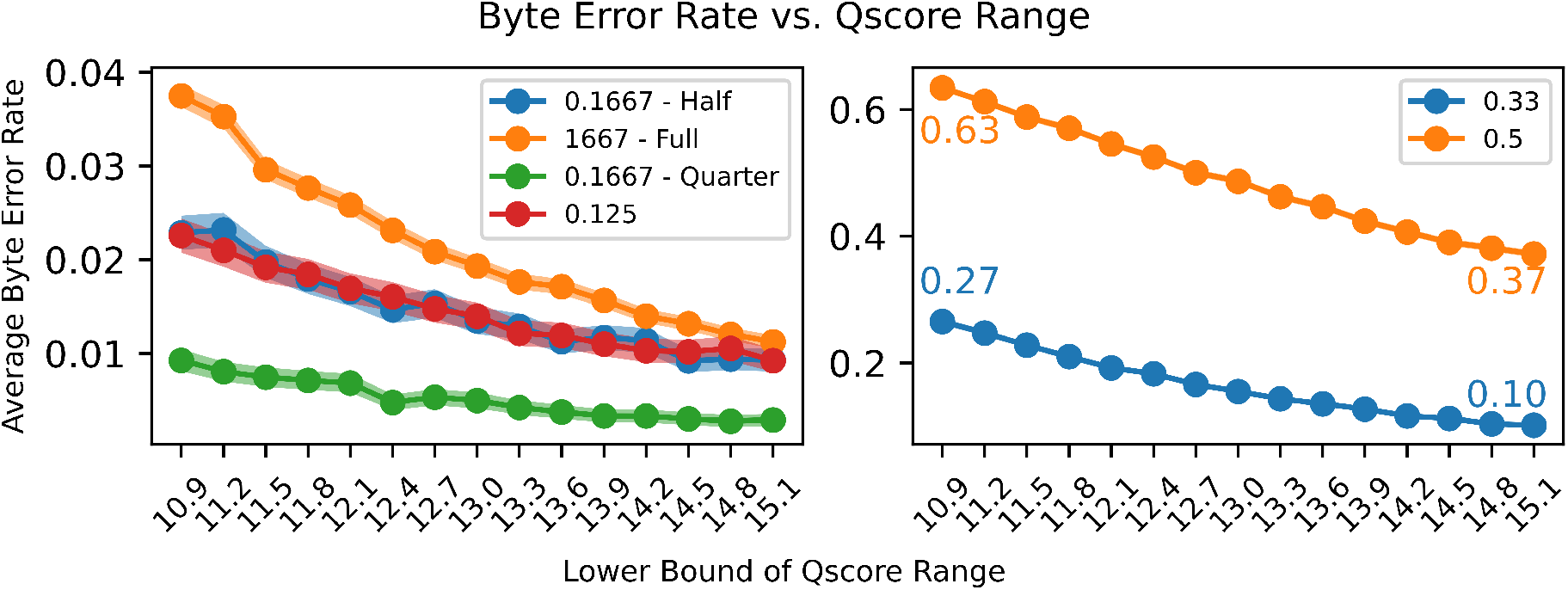
Average byte error rate across all strands for each HEDGES design across a variety of Q score range bins, each with a range of 0.3. Error bands included for each line are calculated for a 95% confidence interval of Supplemental Equation 6. Legend entries represent different the different designs (*ϕ*_HEDGES_ = 0.1667 = 1*/*6), and Quarter, Half, Full, represent the three lengths encoded used for *ϕ*_HEDGES_ = 1*/*6 in the order of increasing length.

## D. Q Score Analysis

To analyze the impact of quality of read provided to the decoder, we construct 15 bins of range 0.3 between Q Scores of 10.9 and 15.1 (Supplemental Figure 16). For each bin, and for all reads of each strand that fall into each bin, we select a 10k random subset for a total of 2.55 million reads. Each point represents 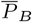 calculated over all positions and strands for a given design.

## E. Description of the Alignment Matrix Algorithm

Here we present pseudo code algorithmic descriptions of our Alignment Matrix algorithm that decodes a convolutional code’s trellis by using CTC scores provided by some machine learning model. Our algorithm relies on manipulating values stored in multi-dimensional arrays. In our notation, we use brackets and indexes to represent retrieving a value from an array. For example, given a three-dimensional array *A*, we access a value at a row *r*, column *c*, and 3-rd dimension *z* by the statement *A*[*r, c, z*]. In scenarios when convenient, we utilize array slicing notation. For example, to generate a 1-dimensional column vector for the same three-dimensional array we write *A*[:, *c, z*] where “:” represents retrieving all row values for a constant column *c* and 3rd dimension *z*. We also utilize multi-dimensional argmax functions that apply in specified dimension of an input array. For example, *B* = **argmax**(*A*, **dim** = 2) outputs a 2-dimensional array with values of the positions of maximum values along the 3rd dimension of *A*. That is, *B*[*r, c*] = argmax_*i*_(*A*[*r, c, i*]).

We break our complete algorithm down into four main sub-algorithm components. The first component is presented in Algorithm 3, which is the top level function to our custom base calling process. Key arguments for this function are the CTC scores (𝒞) that are generated by the machine learning model applied to the raw electrical signal produced by pores, and an array which describes the structure of the convolutional code’s trellis (ℐ). The trellis structure array, ℐ, is 2-dimensional with each row representing a single state in the trellis and each column representing an edge entering each of these states. Each value stored in ℐ is a tuple (*i, v*) which represents a predecessor state *i* with transition value *v*. The main responsibility of this function is to create arrays used to store alignment scoring, and to update those arrays after each step of the trellis. Initially, arrays are created from the viewpoint of transitioning away from each state. That is, we determine the symbols and scores for the edges that transition between step *l* − 1 and *l* of the trellis. Here, we assume that bases are generated by a generic convolutional encoding algorithm (*ConvolutionaGetNextBase*) that generates a base for some convolutional state *h* and transitional value *e*. Outgoing scores from each state are generated by the function *ForwardStep*, which is the core scoring mechanism of our novel algorithm. Then, we rearrange this information via ℐ to create an incoming viewpoint for each state at step *l* such that we can determine the best incoming edge scores and their corresponding bases. Once the best incoming edges are determined, a 2-dimensional array representing the back trace path through the trellis is updated for each state at trellis step *l*. Once all ℒ steps of the trellis complete, *StringFromBacktarce* is used to generate the final basecall from the back trace array.

The scoring function, *ForwardStep* determines scores in 2 phases. The first phase is given in Algorithm 1. Here, an array representing the newly added bases (and inserted blanks) along each state’s outgoing paths, *B*_targets_, is used to determine where symbols repeat. For example, for a given state *h*, transition *e*, and base *G* on edge *e* leaving *h* the vector *B*_targets_[*h, e*, :] represents *b*^′^ − *G*. Here, *b*^′^ is the most recent base that has been aligned for the message currently stored at *h*. Thus, ℱ[:, *h*] is precisely the alignment information stored for base *b*^′^. Knowing where repeats occur is important, as there cannot be any alignment that transitions between the matching bases without passing through a blank symbol. Based on this, we create a masking array *M* which is used to mask out illegal transition scores during the core alignment calculations. Subsequently, all necessary arrays are transferred from CPU memory to a *GPU* device’s memory and threading parameters are constructed for parallel execution of the *ForwardStepKernel* function. We assume *CUDA*-style notation for parameterizing the organization of threads and local (shared) memory. That is, threads are organized in 3-dimensional blocks, and blocks are organized on a 3-dimensional grid. In our algorithm we allocate shared memory for an array A which is designed to hold intermediate alignment scores of single time steps for all symbols whose alignment requires calculation. Finally, *GPU*.*LaunchKernel()* is performed to start execution of the kernel on a GPU device.

The core calculations that determine scores of bases that lie along each edge leaving a trellis state are performed in Algorithm 2. To begin, the array 𝒜 is initialized with values that represent the alignment of symbols at time *T*_*L*_−1, e.g. the time step previous to the lowest time allowed. The array 𝒜 has a value reserved for each base in *B*_targets_ (plus 1 padding base) for the trellis states and edges that execute in the same thread block. We also include a fourth dimension to 𝒜 of length 2 to enable writing the next values of 𝒜 without overwriting the current values that may still be needed by other threads. As the main loop of this algorithm proceeds, 3 values are read from 𝒜 for each symbol that requires alignment calculation. The 3 values represent alignment scores of the previous time step coming from the same symbol, the 1st previous symbol, or the symbol 2 positions away. These values are combined with the raw CTC score for a base *b* at time *t* (𝒞[*t, b*]) to general a final score *S*_final_. *S*_final_ is then used to fill the 𝒜 array position of each base, and a rolling accumulation across time steps of final scores, *R*, is kept for each base. At the end of each loop iteration, the last base scores are written to ℱ_*out*_, and threads in a block are synchronized so that updated 𝒜 values are visible to every thread in the subsequent iteration. As the algorithm terminates, the rolling score *R* is written to *S*_out_ in the thread that represents the final base.

After repeating the process ℒ times of calculating scores and choosing the best incoming edges for each trellis state, Algorithm 4 is executed to build a final base call from chosen paths. For this algorithm, a start state is chosen based on the position of the best score in *S*_current_ of Algorithm 3. Starting at this state, and the last position in the message, the *BT*_index_ array is traversed based on the index stored for each state that points to the best state at the previous position in the message. Along the way, the corresponding positions in *BT*_bases_ are retrieved to build the output basecall. After building a complete message, the string is returned and our algorithm finishes.

### Algorithm 1

Set up function for launching the core GPU kernel threads that calculate alignments in parallel.

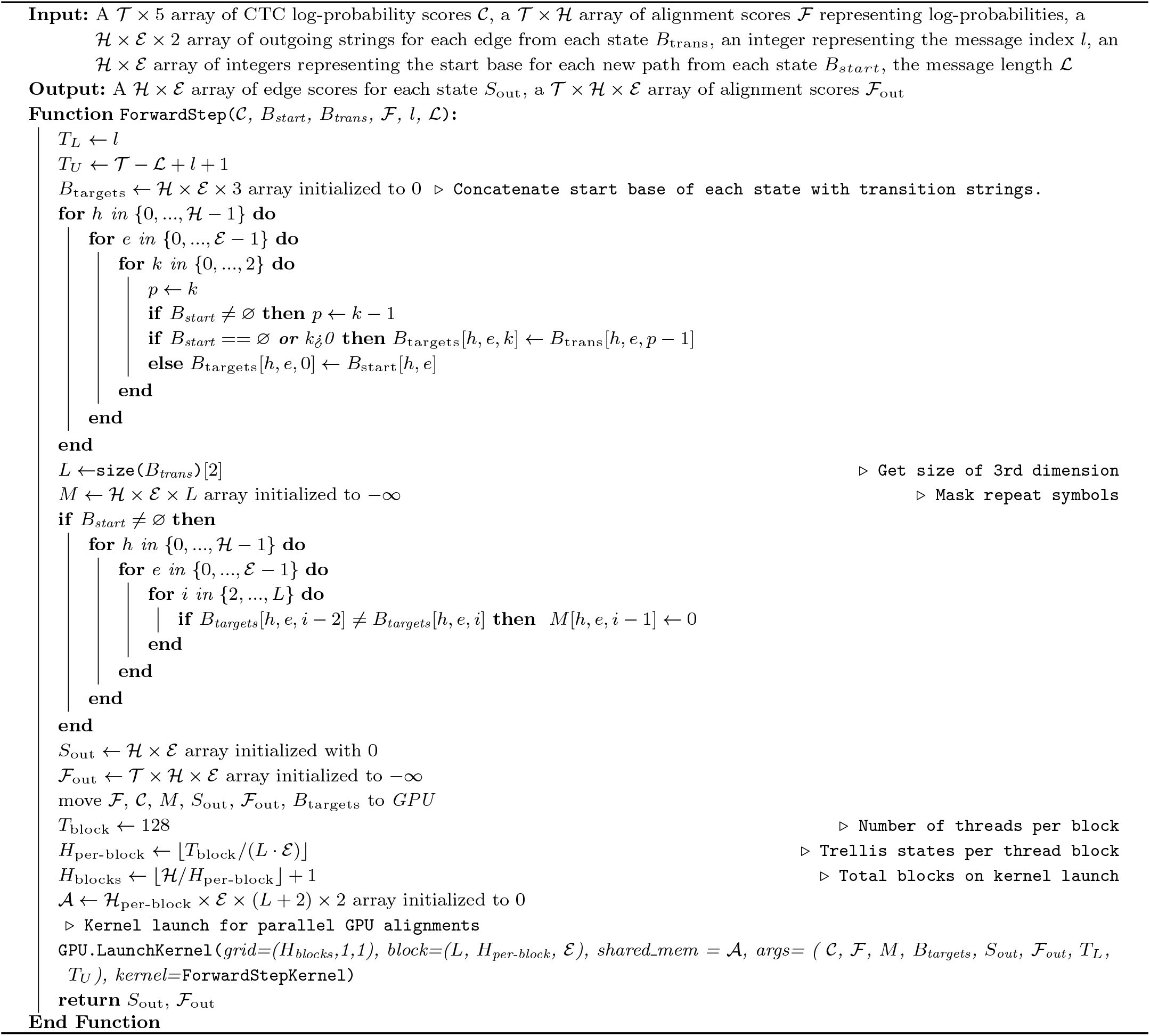

### Algorithm 2

GPU kernel algorithm used to perform the core alignment calculations.

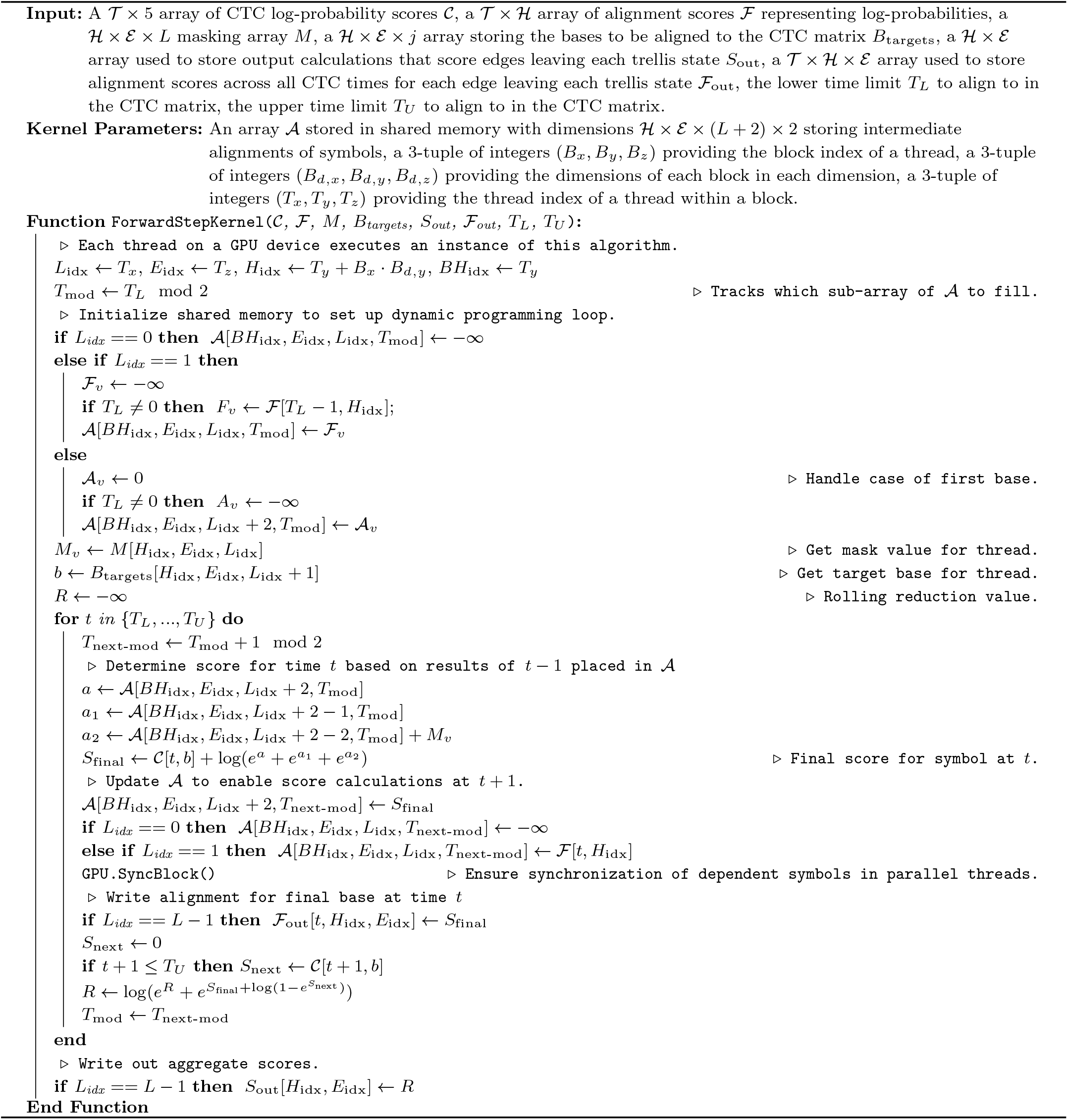

### Algorithm 3

Main basecalling function that determines a sequence of bases from an input trellis structure and CTC scoring of bases across a read’s electrical signal.

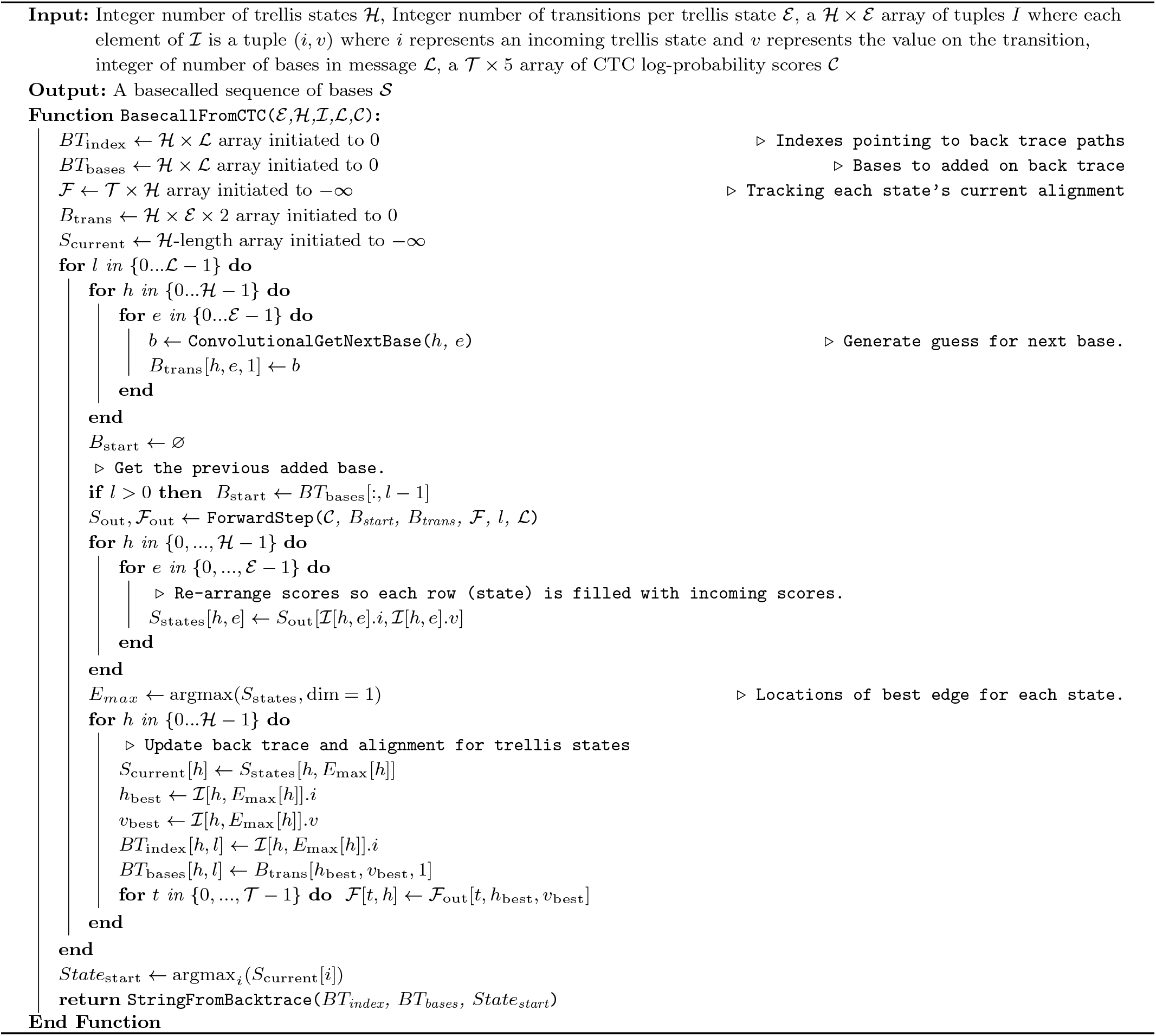

### Algorithm 4

Algorithm that derives an output base sequence based on the computed backtrace matrices.

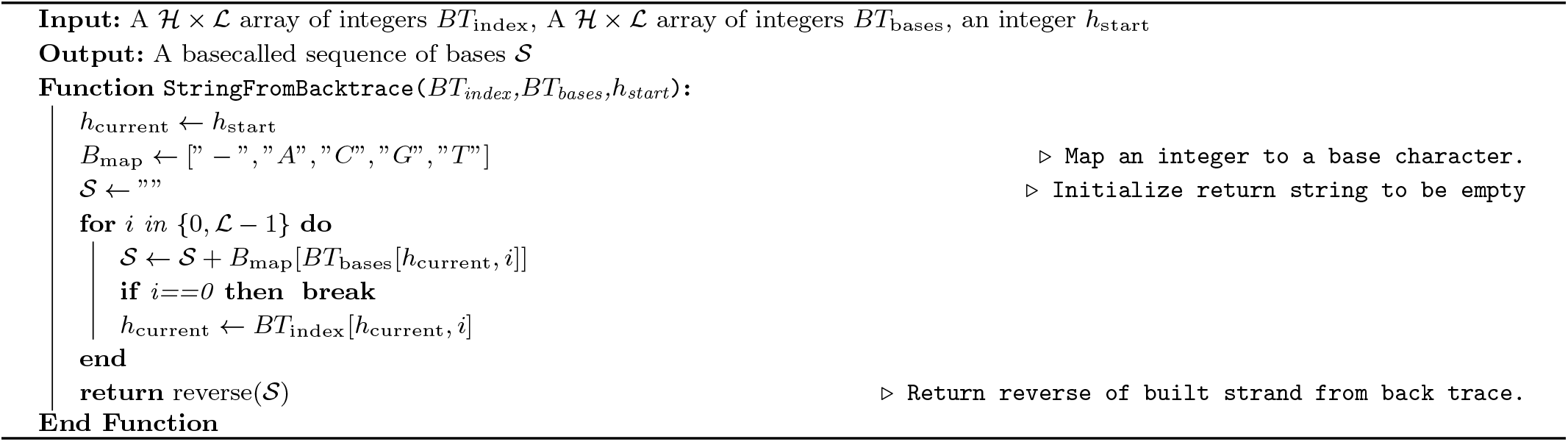

**Table 1.**
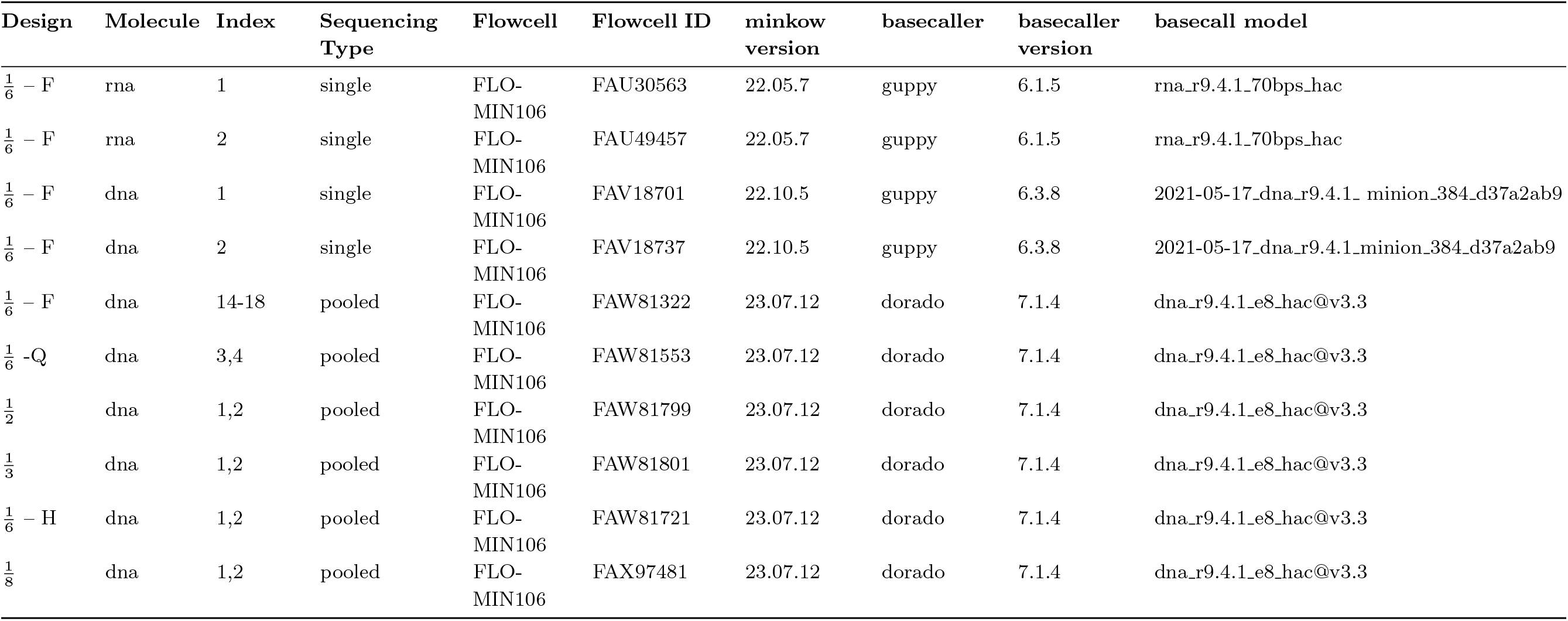
Sequencing run and basecaller specification for the nanopore sequencing of each encoded strand. The designs labeled with F (full)/H (half)/Q (quarter) refer to the three sizes of the 1*/*6 HEDGES rates code that was synthesized in decreasing length order.

## References

Chandak, S. et al. (2020) Overcoming High Nanopore Basecaller Error Rates for DNA Storage via Basecaller-Decoder Integration and Convolutional Codes, ICASSP 2020 -2020 IEEE International Conference on Acoustics, Speech and Signal Processing (ICASSP), pages 8822–8826. ISSN: 2379-190X.

Chen, W. et al. (2021) An artificial chromosome for data storage, National Science Review, 8(5)nwab028.

Choi, Y. et al. (2019) High information capacity DNA-based data storage with augmented encoding characters using degenerate bases, Scientific Reports, 9(1)1–7.

Graves, A. et al. (2006) Connectionist temporal classification: labelling unsegmented sequence data with recurrent neural networks. In Proceedings of the 23rd international conference on Machine learning, ICML ‘06, pages 369–376, New York, NY, USA Association for Computing Machinery. ISBN 978-1-59593-383-6.

Kovaka, S. et al. (2021) Targeted nanopore sequencing by real-time mapping of raw electrical signal with UNCALLED, Nature Biotechnology, 39(4)431–441.

Kürzinger, L. et al. (2020) CTC-Segmentation of Large Corpora for German End-to-End Speech Recognition. In Karpov, A.and Potapova, R., editors, Speech and Computer, Lecture Notes in Computer Science, pages 267–278, Cham Springer International Publishing. ISBN 978-3-030-60276-5.

Neumann, D., Reddy, A. S. N., and Ben-Hur, A. (2022) RODAN: a fully convolutional architecture for basecalling nanopore RNA sequencing data, BMC Bioinformatics, 23(1)142.

Nguyen, B. H. et al. (2021) Scaling DNA data storage with nanoscale electrode wells, Science Advances, 7(48)eabi6714.

Organick, L. et al. (2018) Random access in large-scale DNA data storage, Nature Biotechnology, 36(3)242–248.

Pagés-Gallego, M. and de Ridder, J. (2023) Comprehensive benchmark and architectural analysis of deep learning models for nanopore sequencing basecalling, Genome Biology, 24(1)71.

Press, W. H. et al. (2020) HEDGES error-correcting code for DNA storage corrects indels and allows sequence constraints, Proceedings of the National Academy of Sciences, 117(31)18489–18496.

Scheidl, H., Fiel, S., and Sablatnig, R. (2018) Word Beam Search: A Connectionist Temporal Classification Decoding Algorithm. In 2018 16th International Conference on Frontiers in Handwriting Recognition (ICFHR), pages 253–258.

Tomek, K. J. et al. (2019) Driving the Scalability of DNA-Based Information Storage Systems, ACS Synthetic Biology, 8(6)1241– 1248.

Volkel, K. D. et al. (2023) FrameD: framework for DNA-based data storage design, verification, and validation, Bioinformatics, 39(10) btad572.

Wang, Y. et al. (2021) Nanopore sequencing technology, bioinformatics and applications, Nature Biotechnology, 39(11)1348–1365.

Wick, R. R., Judd, L. M., and Holt, K. E. (2019) Performance of neural network basecalling tools for Oxford Nanopore sequencing, Genome Biology, 20(1)129.

Yazdi, S. M. H. T., Gabrys, R., and Milenkovic, O. (2017) Portable and Error-Free DNA-Based Data Storage, Scientific Reports, 7(1) 5011.

